# Molecular basis of N_2_ fixation in a hyperthermophilic archaeon

**DOI:** 10.1101/2025.10.10.681579

**Authors:** Nevena Maslać, Mustafa Rasim Törer, Pauline Bolte, Tristan Wagner

## Abstract

Exploring the natural diversity of phylogenetically distant nitrogenases is crucial for gaining new insights into the mechanism of atmospheric N_2_ fixation and unlocking biotechnological developments in sustainable ammonia production. Here, we investigated the N_2_-fixing system of *Methanocaldococcus infernus*, a deep-sea hyperthermophilic archaeon growing diazotrophically at 92 °C. This natively isolated nitrogenase has a melting temperature close to the water boiling point, with an extrapolated specific activity superior to mesophilic homologues. The crystal structures obtained at near-atomic resolution present the most simplified known nitrogenase, harbouring strategic hot spots for thermostability. It combines the structural traits of all three known nitrogenase isoforms, reinforcing the postulate that the archaeal enzyme predates modern versions. In contrast to structural homologues, the electron-transferring metallocofactor “P-cluster” is trapped in a rare state awaiting electron delivery, providing a detailed picture of the physiological state. The active site, harbouring the FeMo-cofactor catalyst, exhibits a mixture of the resting and “turnover” states, previously described solely in the bacterial vanadium and iron-only nitrogenases. Therefore, these results unify a mechanistic principle of all nitrogenases and highlight the advantages of the hyperthermostable nature of the archaeal enzyme, opening new avenues for further understanding of how nature splits the N_2_ triple bond.

## Introduction

Performed exclusively by diazotrophic microbes, biological N_2_-fixation provides half of the nitrogen to the living organisms of our planet (1–3). Alongside a minor contribution from geochemical discharge through lightning, diazotrophy is the only known natural reaction capable of breaking the triple bond of atmospheric N_2_ to generate the bioavailable fertiliser and biofuel ammonia (NH_3_) under ambient pressure and temperature (4). In comparison, the industrial Haber-Bosch process requires extreme conditions (200 bar and 400-500 °C) and generates 1–2% of the global CO_2_ emissions (5–8). Greener alternatives for decarbonising NH_3_ production could be inspired by biological systems, for instance, to transform eukaryotes into diazotrophs (9, 10). For this, a deep understanding of the biological N_2_ fixation mechanism, which remains one of the most complex unanswered questions in bioinorganic catalysis, is necessary and must be addressed.

The sole enzymatic system capable of fixing N_2_ is the ATP-dependent nitrogenase, composed of NifH and NifDK. NifH contains a [4Fe-4S]-cluster capable of delivering an electron upon ATP hydrolysis to NifDK (3, 11). Electrons are ultimately transferred to the FeMo-cofactor bound on NifD (FeMoco, one of the most complex enzymatic metallocofactors known in living systems) through the P-cluster. The P-cluster, harboured at the NifDK interface, has been proposed to oscillate between a fully reduced P^N^ and the electron-depleted P^1+^ state (following the accepted deficit-spending mechanism, Supplementary Fig. 1a) (12–14). There are two more isoforms of nitrogenase in which the FeMoco is replaced by a FeV-cofactor (Vnf system) or a FeFe-cofactor (Anf system, Supplementary Fig. 1a)(15, 16).

Spanning more than 60 years, research integrating structural, spectroscopic, and biochemical data has deepened our understanding of the complexity underlying biological N₂ reduction. Yet, the field remains largely shaped by studies on a narrow set of diazotrophic organisms. Most of the knowledge stems from *Azotobacter vinelandii*, whose disproportionately large set of *nif* genes may hinder our perception. Phylogenetically distant diazotrophs, on the other hand, with distinct lifestyles and metabolic requirements, often rely on a minimal set of genes for N₂ fixation. Such a shift in perspective not only fortifies ongoing synthetic biology initiatives (9, 17) but offers the opportunity to clarify the mechanistic principles of N₂ fixation by distinguishing conserved strategies and revealing unanticipated modes of adaptation within nitrogenase catalysis.

In this work, we explore the distant nitrogenase from marine *Methanococcales*, which have been established by phylogenetic studies as the progenitor of modern nitrogenase systems (18–22). Diazotrophic *Methanoccocales* have been highlighted for their environmental importance as primary N_2_-fixers in some ecological niches (23–25), successful transplantation of part of their nitrogenase system in plants (26, 27), differences in the regulation of their nitrogen-assimilation node (28–32), and for their ability to fix N_2_ at the highest temperature recorded so far of 92 °C (33). By studying the nitrogenase from *Methanocaldococcus infernus*, we found molecular adaptive traits for hyperthermostability and shared structural characteristics with all three known nitrogenase isoforms, providing new insights into the N_2_-fixation mechanisms.

### Native isolation of a hyperthermostable nitrogenase

With its minimal *nif* operon (Supplementary Fig 1b), *M. infernus* presents remarkable behaviour during diazotrophic lifestyle with a doubling time of 2 hours under complete autotrophy (Fig. 1a), at a maximum temperature of 91.7 °C, withstanding temperatures as high as 93 °C, though viability is lost after 24h (Supplementary Fig. 2a). Despite reaching high cell density, the requirement for both nitrogen sources differs: 20 mmol of N_2_ is required for optimal diazotrophic growth in contrast to only 0.22 mmol for NH_4_Cl (Fig. 1a, Supplementary Fig. 2b). This behaviour might come from a low solubilisation of N_2_ in the medium at 75 °C, a difference of affinity between the nitrogenase and the glutamine synthetase (i.e., NH_3_-fixation (31)), or could underline a competitive inhibitory effect of H_2_ versus N_2_ as reported for *A. vinelandii* nitrogenase (34).

**Fig. 1.**
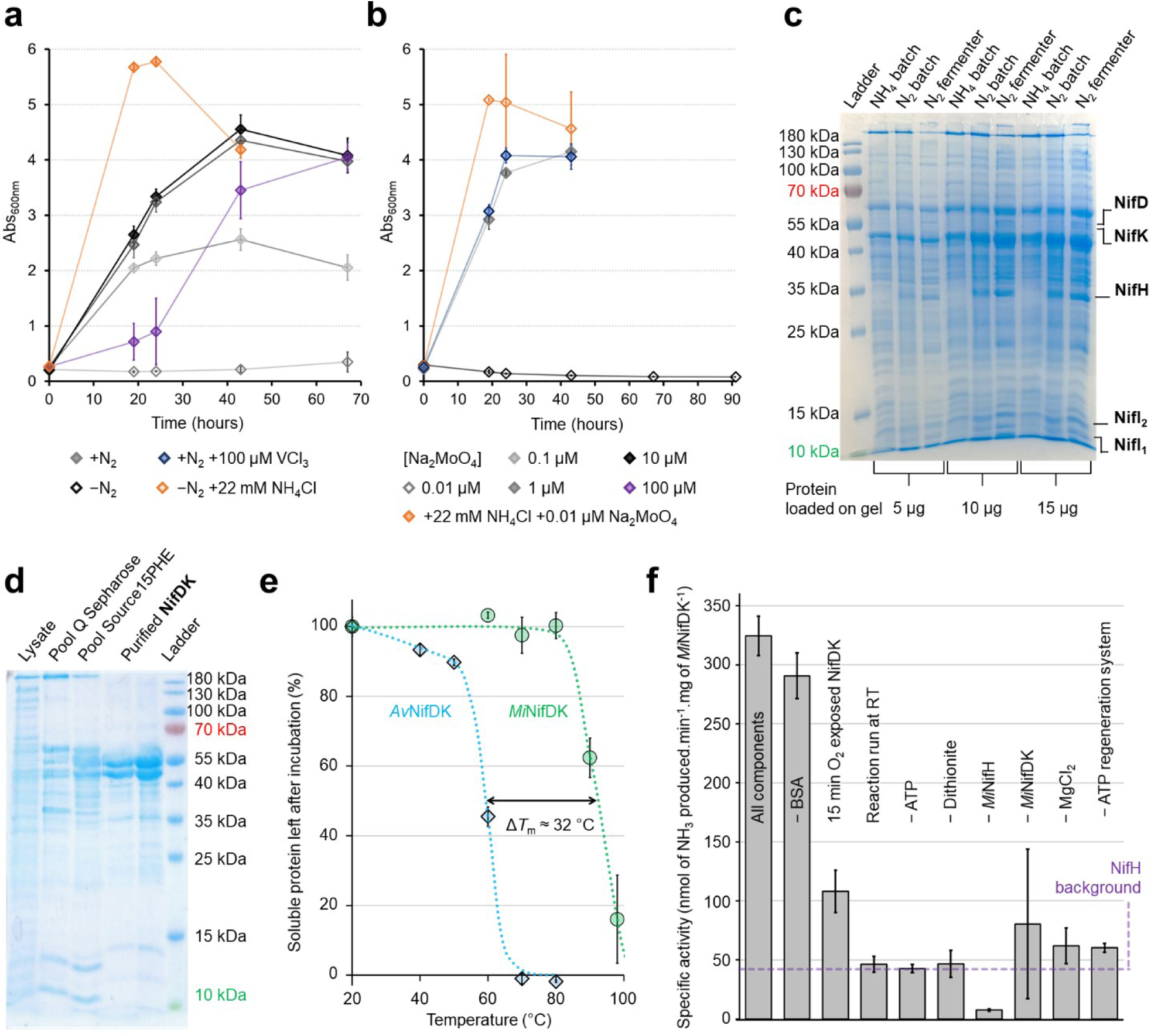
Isolation of an active hyperthermostable nitrogenase from the diazotroph *M. infernus*. **a**, Diazotrophic growth *M. infernus* on 10 µM Na_2_MoO_4_ and **b,** its molybdenum dependency. All tested conditions contain 0.5 bar of N_2_, 100 µM Na_2_WO_4_ and were grown at 75 °C. **c**, *M. infernus* cell extracts grown on NH_4_Cl versus N_2_ in batch culture or in a fermenter showing a natural overexpression of the *nif* system. In a-b, a final quantity of 0.22 mmol NH_4_Cl was added. **d,** Native purification of *Mi*NifDK. To separate the regulator *Mi*NifI_1,2_ from NifDK, 2-oxoglutarate and MgATP were added to the Source^TM^ 15PHE pool before further separation by anionic exchange chromatography. 3 µg of proteins were loaded on each well, the NifDK sample close to the ladder being a fraction slightly more contaminated. **e,** Thermostability assay of *Av*NifDK and *Mi*NifDK, presenting an estimated *T*_m_ of 60.1 °C and 92.3 °C, respectively. Data from *Av*NifDK comes from Maslać et al. 2024 (41). Dashed lines come from manual fits to illustrate the trend. **f,** Activity assay of *Mi*NifDKH performed at 50 °C under 100% N_2_ at pH 7.0 for 30 min. Experiments in **a, b** and **f** have been performed in triplicate, while experiments in panel **e** were performed in duplicate.

Diazotrophic growth of *M. infernus* is molybdenum dependent (Fig. 1b), insensitive to VCl_3_ addition (Fig. 1a), and exhibits an outstanding tolerance to tungstate: up to 1,000 times higher than what has been observed in *A. vinelandii* or *Methanococcus maripaludis*, a mesophilic *Methanococcales* (35, 36). The latter trait, also reported on the thermophile *Methanothermococcus thermolithotrophicus* (29), might be an adaptation to the environmental conditions. Despite a closer sequence homology to a VFeco-containing nitrogenase (*Av*VnfDK, Supplementary Fig. 1b), the molybdenum-dependent diazotrophic growth suggests the utilisation of a FeMoco.

To understand the true nature of *M. infernus* nitrogenase, rather than relying on recombinant systems, we studied the enzyme directly isolated from diazotrophically grown cells expressing naturally high quantities of the N_2_-fixation machinery (37, 38) (Fig. 1c). During the purification, *Mi*NifDK coeluted with *Mi*NifI_1,2_, a regulatory complex belonging to the P_II_-family proteins detecting cellular energy (i.e., ATP/ADP/AMP ratio) and cellular nitrogen availability (i.e., 2-oxoglutarate/glutamate/glutamine ratio)(28, 32, 39). *Mi*NifI_1,2_ binding to *Mi*NifDK is most likely due to the decrease of MgATP and 2-oxoglutarate concentration during cell lysis (Fig. 1d)(40). As expected, addition of these ligands release *Mi*NifI_1,2_ inhibition and allows *Mi*NifDK separation and isolation (Supplementary Fig. 2c, Fig. 1d).

The melting temperature (*T*_m_) of *Mi*NifDK was estimated to be 92.3 °C, which is roughly 32 °C higher than that of *Av*NifDK reported previously (41) (60.1 °C, Fig. 1e). This is consistent with the diazotrophic growth at 91.7 °C and the high thermostability recorded from the recombinantly produced *Mi*NifH (Supplementary Fig. 2d) with a *T*_m_ of 82.5 °C (41). Unlike nitrogenases derived from mesophilic organisms (i.e., *A. vinelandii*), NH_3_ production is undetectable when *M. infernus* whole nitrogenase system is assayed at room temperature (Fig. 1f), reflecting adaptations to contrasting ecological niches and hyperthermophilic adaptation. When the temperature is increased to 50 °C, the maximal temperature allowed by the ATP-regeneration system, a specific activity of 280 ± 18.2 nmol of NH_3_ produced.min^-1^.mg^-1^ was measured (Fig. 1f). Assuming that the activity follows the Arrhenius law, then this extrapolation would lead to an activity of 2,250 nmol of NH_3_ produced.min^-1^.mg of *Mi*NifDK^-1^ at 80 °C, which would be the highest nitrogenase activity ever measured (≈ 500-900 nmol of NH_3_ produced.min^-1^.mg^-1^ for native *Av*NifDK (42–44)). The temperature-dependent activity suggests that the enthalpic stabilisation that keeps the structure rigid also prevents the local flexibility needed for substrate turnover, stalling the enzyme.

### General structural features

The strong reductant dithionite (S_2_O_4_^2-^, DT) is routinely included in nitrogenase purifications, although recent studies have suggested that dithionite may be a non-innocent reductant for metalloproteins (45). Thus, here, the relatively mild reductant dithiothreitol was added during anaerobic purification and crystallisation, leading to a partially oxidised preparation (Supplementary Fig. 2e). Two crystalline forms named A (monoclinic) and B (triclinic) were obtained yielding high-quality crystals, which allowed the refinement of the structures to a resolution of 1.37 Å and 1.21 Å, respectively (Supplementary Table 1).

The overall structure presents the canonical NifD_2_K_2_ assembly in which dimerisation occurs via NifK (Fig. 2a). As observed by sequence identity conservation (Supplementary Fig. 1b) and phylogeny analyses (21), *Mi*NifDK is not structurally more similar to any particular nitrogenase homologue (Fig. 2a, Supplementary Fig. 1b), limiting its assignment to a Nif, Vnf or Anf system. To resolve this uncertainty and to confirm whether this archaeal enzyme belongs to a Nif system, X-ray diffraction anomalous-based metal identification was conducted. The molybdenum signature was unambiguously confirmed by comparing the anomalous signal from datasets collected at energies before and at the peak of fluorescence of Mo (Fig. 2a and Supplementary Table 1). A similar experiment was conducted on the Fe K-edge, providing high confidence in the Fe signature for the FeMoco and the P-cluster (site numbers 1 and 2 on Fig. 2a). The absence of signal excluded the presence of an Fe atom at the NifK/NifK′ interface (site number 3 on Fig. 2a, also called the 16^th^ Fe site (44, 46), which adequately fits a Mg^2+^ as modelled in *Av*VnfDK and *Av*AnfDK (42, 47).

**Fig. 2.**
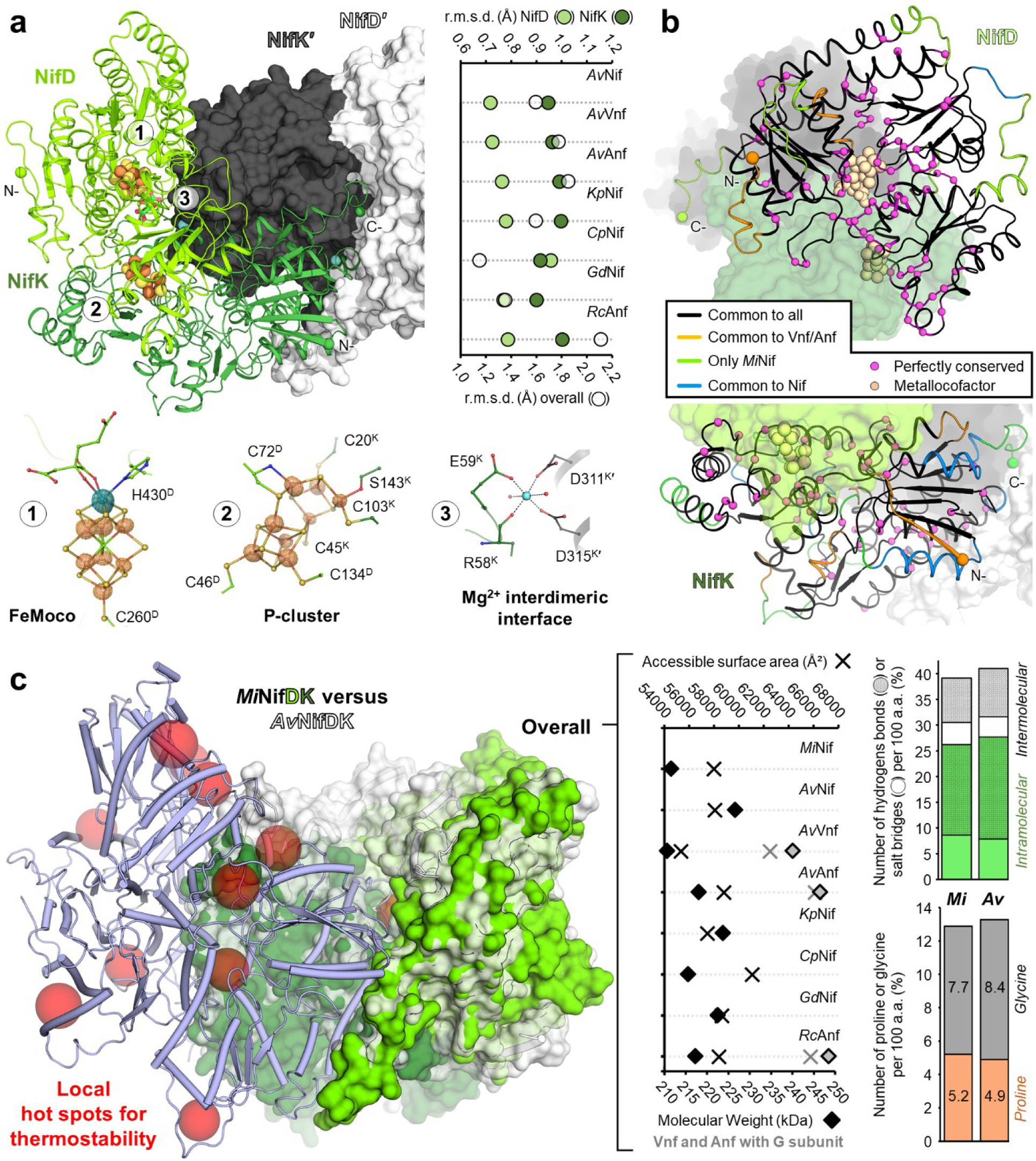
Structural conservation and thermostability features of *Mi*NifDK. **a**, *Mi*NifDK overall view with one NifDK in cartoon and the other in surface. Metallocofactors are displayed as balls and sticks with a close-up (bottom panel). Anomalous difference maps for molybdenum (blue mesh/surface) and iron (orange mesh/surface) are contoured at 6 and 8 σ, respectively. Right graph represents the root mean square deviations between *Mi*NifD, *Mi*NifK or *Mi*NifD_2_K_2_ and respective structural homologues. **b**, Secondary structure conservation between *Mi*NifDK and homologues after manual inspection. *Mi*NifD (top) and *Mi*NifK (bottom) are shown as cartoons with the other subunits shown as surface and coloured as in **a,** with perfectly conserved residues represented as spheres. **c,** Superposition of *Mi*NifDK (green) and *Av*NifDK (transparent white surface and white cartoon). The second *Mi*NifDK is depicted in a purple cartoon, with local hot spots indicated by red spheres. The right graphs compare surface accessibility, molecular weight, the number of hydrogen bonds/salt bridges, and the Gly/Pro ratio between *Mi*NifDK and *Av*NifDK.

*Mi*NifDK is a FeMoco-containing nitrogenase, but it shares structural features with both Vnf and Anf systems (Fig. 2b). Among shared structural traits are the N-terminal parts of *Mi*NifD and *Mi*NifK, some loops and alpha-helices, and short sections, while structural similarity with Nif is rather limited to a loop in NifD and a few sections in NifK. The C-terminal parts and some peripheral loops of NifD and NifK are unique to *M. infernus* (Fig. 2b). The leftover core conserved across all structural homologues (representing 84.6 and 74 % of the residues in *Mi*NifD and *Mi*NifK, respectively) contains identical residues maintaining structural integrity (glycine and proline), binding the metallocofactor and involved in the catalysis. Additionally, conserved positions are numerous at dimeric interfaces, probably to maintain important interactions for the quaternary structure.

Deep structural comparison with *Av*NifDK provides a rational basis to explain the hyperthermostability of *Mi*NifDK (Fig. 1e). The overall structure of *Mi*NifDK represents the most simplified scaffold compared to Nif bacterial homologues (Fig. 2c and Supplementary Figs. 3-5). This results in a reduced overall size, comparable with VnfDK without the VnfG subunit, but does not decrease the surface exposed to the solvent (e.g., no reduction in unstable loops or domains). General traits expected to increase overall stability (i.e., additional inter/intramolecular hydrogen bonding network and salt bridges, reduction of glycine and increase of proline, clusters of hydrophobic residues) are only observable to some extent between *Mi*NifDK and *Av*NifDK (Fig. 2c, Supplementary Fig. 6). Hence, we propose that the hyperthermostability is the consequence of a rearrangement of salt bridges/hydrogen bonds and hydrophobic clustering at strategic points. These “hot spots” for structural stability would reinforce the local contacts, while the reallocation of proline and glycine would allow the protein to balance rigidity and flexibility where it is needed (Fig. 2c, Supplementary Figs. 6b and 7). Two of these points are located on the αIII domain (composed of residues 2-32 and 334-430 in *Mi*NifD based on (48)), known for its high intrinsic flexibility and participating in FeMoco delivery and catalysis (49–51). It is worth noting that the hot spots are mostly located on the surface, distant from the FeMoco and P-cluster sites, suggesting that the active site and electron transfer path are already highly stabilised or depend on intrinsic flexibility for optimal catalysis.

### The electron transfer path

A comparison of the electron transfer path within *Mi*NifDK and the structural homologues from *A. vinelandii* shows a few divergences in the vicinity of the P-cluster, with some positions in the outer shell shared with *Av*VnfDK/*Av*AnfDK and not *Av*NifDK (Fig. 3a). Most importantly, the relay involved in the electron transfer from the P-cluster to the FeMoco is identical to that of the Nif system instead of Vnf/Anf (i.e., the Tyr48^K^/A49^D^ pair in *Mi*NifDK instead of the Phe59^K^/Cys52^D^ pair in *Av*Vnf and Phe48^K^/Cys52^D^ in *Av*Anf)(42, 47). As expected, the P-cluster-anchoring cysteines and a few residues pointing towards the NifH binding surface are perfectly conserved.

**Fig. 3.**
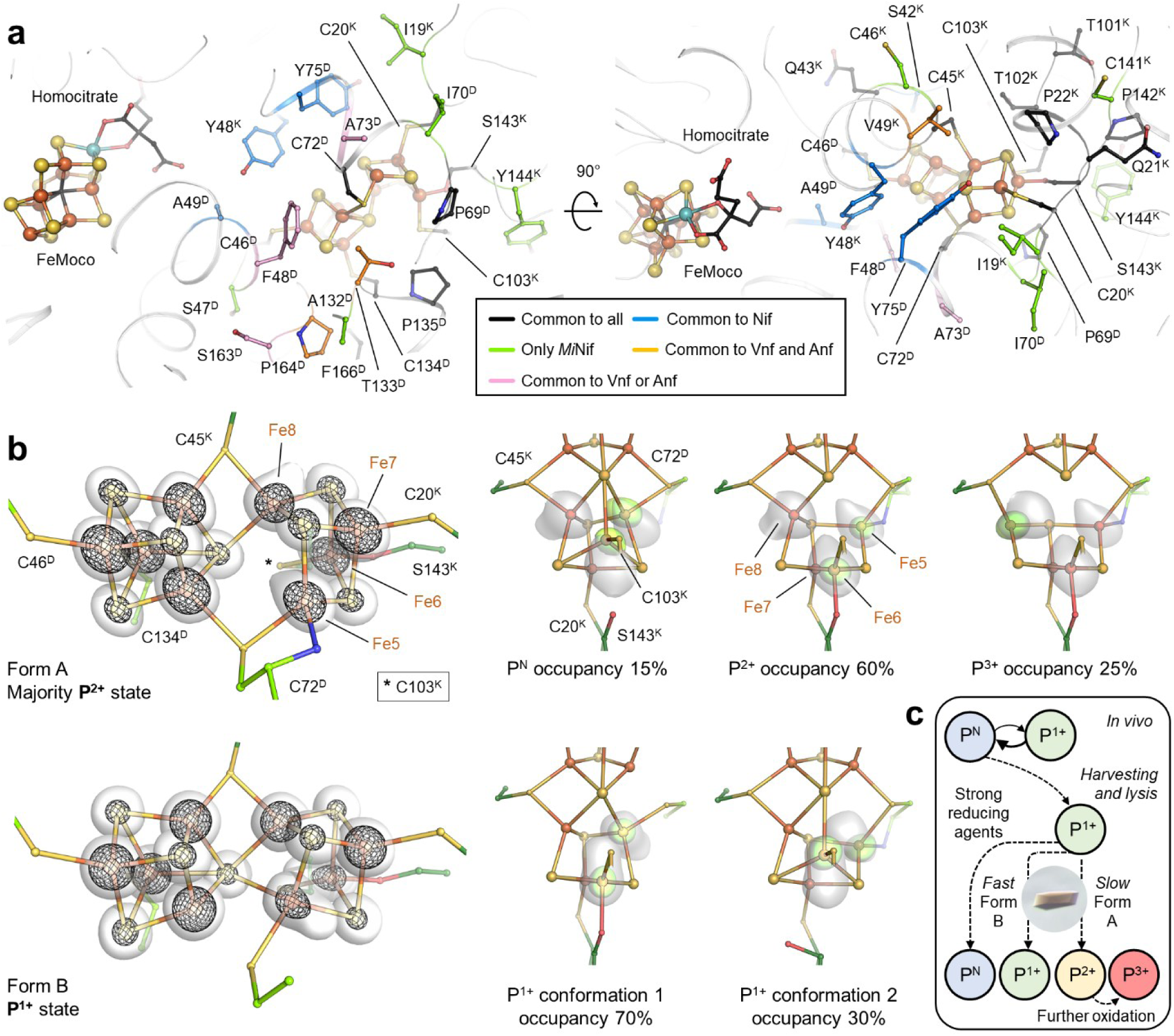
P-cluster site. **a**, Conservation of the P-cluster surroundings in *Mi*NifDK and structural homologues. The protein backbone is shown as a cartoon, with residues which are part of the 1^st^ and 2^nd^ spheres displayed as sticks. Metallocofactors are displayed as spheres and sticks. A colour coding of carbon shows the conservation among *Av*NifDK, *Av*VnfDK and *Av*AnfDK. **b**, Close up of the P-cluster in the form A (top) and form B (bottom) with Fe and S from the metallocofactor displayed as spheres and residues binding the P-cluster as sticks. 2*F*_o_-*F*_c_ maps are overlaid as a transparent surface (3.5-σ) and black mesh (8-σ). Right panels illustrate the interpretation of the mixture of different states deconvoluted as a P^N^, P^2+^ and P^3+^ for the form A (top) and two alternative conformations of the P^1+^ state for the form B (bottom). The 2*F*_o_−*F*_c_ contoured to 2 σ is presented as a transparent white surface. An omit map was generated for Fe5, Fe6 or Fe8 for the corresponding P-cluster, and the resulting *F*_o_−*F*_c_ maps are presented as a green transparent surface contoured to 7-σ, 30-σ, 10-σ, 32-σ, and 11-σ for P^N^, P^2+^, P^3+^, P^1+^ alternative conformer A, and P^1+^ alternative conformer B, respectively. **c**, Scheme illustrating the proposed scenario for how the different isolated states of the P-cluster were obtained. Here, due to the absence of a reducing agent and fast crystallisation, the crystalline form B kept the P^1+^ state, while the crystalline form A has probably been progressively oxidised to P^2+^ and P^3+^ because of crystallisation delays.

We inspected the redox state(s) of the P-cluster by analysing its coordination. The form A mainly exhibits the P^2+^ state modelled with an occupancy of 60%. This oxidised state is recognisable by the Fe6 and Fe5 respectively bound to the hydroxyl-Ser143^K^ and backbone amide of Cys72^D^ main chain, and a bond loss with the central sulfur (Fig. 3b, Supplementary Fig. 8a)(12). The remaining 40% are composed of a mixture of two other states. The second conformer is typical for a P^N^ state, with electron densities for Fe6 and Fe5 disengaged from the protein ligands, bound to the central sulfur. As-isolated *Av*NifDK purified in the absence of reducing agents has been reported to only contain the P^2+^ state (52), however, a similar mixture of P^N^ and P^2+^ was seen in other systems (53, 54). The third conformer comes from an electron density deviation around the Fe8, causing a bond breakage with the central sulfur (referred to as P^3+^, Fig. 3b). The same displacement was observed in the P^2+^ state of the nitrogenase from *Gluconocetabacter diazotrophicus* (*Gd*NifDK), where Fe8 is coordinated by Tyr98^K^ (55), being replaced by Phe99^K^ in *Av*NifDK and Val49^K^ in *Mi*NifDK (Supplementary Fig. 8b). This displacement is assumed to counterbalance the loss of the Fe6-Ser coordination due to serine substitution to alanine in *Gd*NifDK, and was proven by exchanging the pair Phe99^K^/Ser188^K^ to Tyr99^K^/Ala188^K^ in *Av*NifDK. In *Mi*NifDK, Fe8 displacement is stabilised by a water molecule placed at the same position as the hydroxy group of the Tyr98^K^ in *Gd*NifDK (Supplementary Fig. 8b). We propose that this state implies a further oxidation of the P-cluster in a P^3+^, and we suspect its preservation to be due to the high enzyme stability caging the oxidised cluster and preventing its collapse.

In contrast to form A, which mostly contains the P^2+^ state, form B contains the P^1+^ state in which Fe6 is bound to Ser143^K^ and Fe5 is bound to the central sulfur (Fig. 3b, Supplementary Fig. 8a). It is a very rare state difficult to trap without redox-mediating agents (56, 57). Elongated electron densities emanating from Fe6 and Fe5 suggest an alternative conformation of the P^1+^ state with an estimated occupancy of 30%. In this conformer, Fe5 is now bound to the amide backbone of Cys72^D^ and Fe6 is bound to the central sulfur. Residues in the surroundings of the P-cluster, including the electron relay, are perfectly superposable between both forms, suggesting that the P^1+^ state does not promote drastic conformational changes that would impact the FeMoco centre, such as internal cross-communication.

The different observed redox states between form A and B are rationalised as follows: (*i*) The crystalline forms might have selected a *Mi*NifDK population trapped in a particular state. An explanation challenged by the fact that both structures are superimposable (overall root mean square deviation of 0.226 Å for 1,723 aligned Cα). (*ii*) Form A comes from a low-concentrated sample with crystals appearing after a month, compared to the crystals of form B, which were obtained from a high-concentrated sample and were immediately flash frozen after appearing within a few days. (Fig. 3c). This explanation implies that during harvesting and lysis, the extracted *Mi*NifDK harboured in majority a P^1+^ state. This state was stabilised over the purification and fast crystallisation by the intrinsic enzyme rigidity and the mild reductant dithiothreitol without causing its reduction to P^N^ in contrast to strong reducing agents such as dithionite (Supplementary Fig. 2e)(12). The crystallisation delay in form A would have provoked a further uncontrolled oxidation event (e.g., partial O_2_ contamination in the glove box), leading to the appearance of the P^2+^ and P^3+^. The very low 15% occupancy of the P^N^ in the form A might be due to X-ray radiation or a mixture of P^1+^ states. Altogether, the crystal structures show the overall spectrum of states of the P-cluster critical to understand the enzyme mechanism, and further studies will document this landscape of redox states with the implication on the FeMoco site.

### A FeMoco site captured in resting and “turnover” states underscores common catalytic principles of nitrogenases

The catalytic chamber residues are largely identical across all isoforms, with *Mi*NifDK specific variations occurring at the bulky Met216^D^ pointing to the molybdenum and Gln462^K^′, Tyr51^K^, Asn428^D^ slightly readjusting the water network close to the homocitrate (Fig. 4a-b and Supplementary Fig. 9a). The hydrogen bonding network made of S172^D^, H175^D^, R262^D^, S263^D^, Y266^D^, and H367^D^ is perfectly conserved and might be involved in proton delivery, a controversial topic (58, 59). The identity of the central ligand of the FeMoco is proposed to be a carbide atom based on the 1.21-Å structure, corroborating that the FeMoco is chemically identical to that of *Av*Nif (Supplementary Fig. 9b). A superposition of the metallocofactors harboured in all *A. vinelandii* isoforms confirmed the almost identical position of Fe and S atoms, as well as the homocitrate between *Mi*NifDK FeMoco and *Av*NifDK FeMoco with slight deviations within *Av*VnfDK and *Av*AnfDK (Supplementary Fig. 9c).

**Fig. 4.**
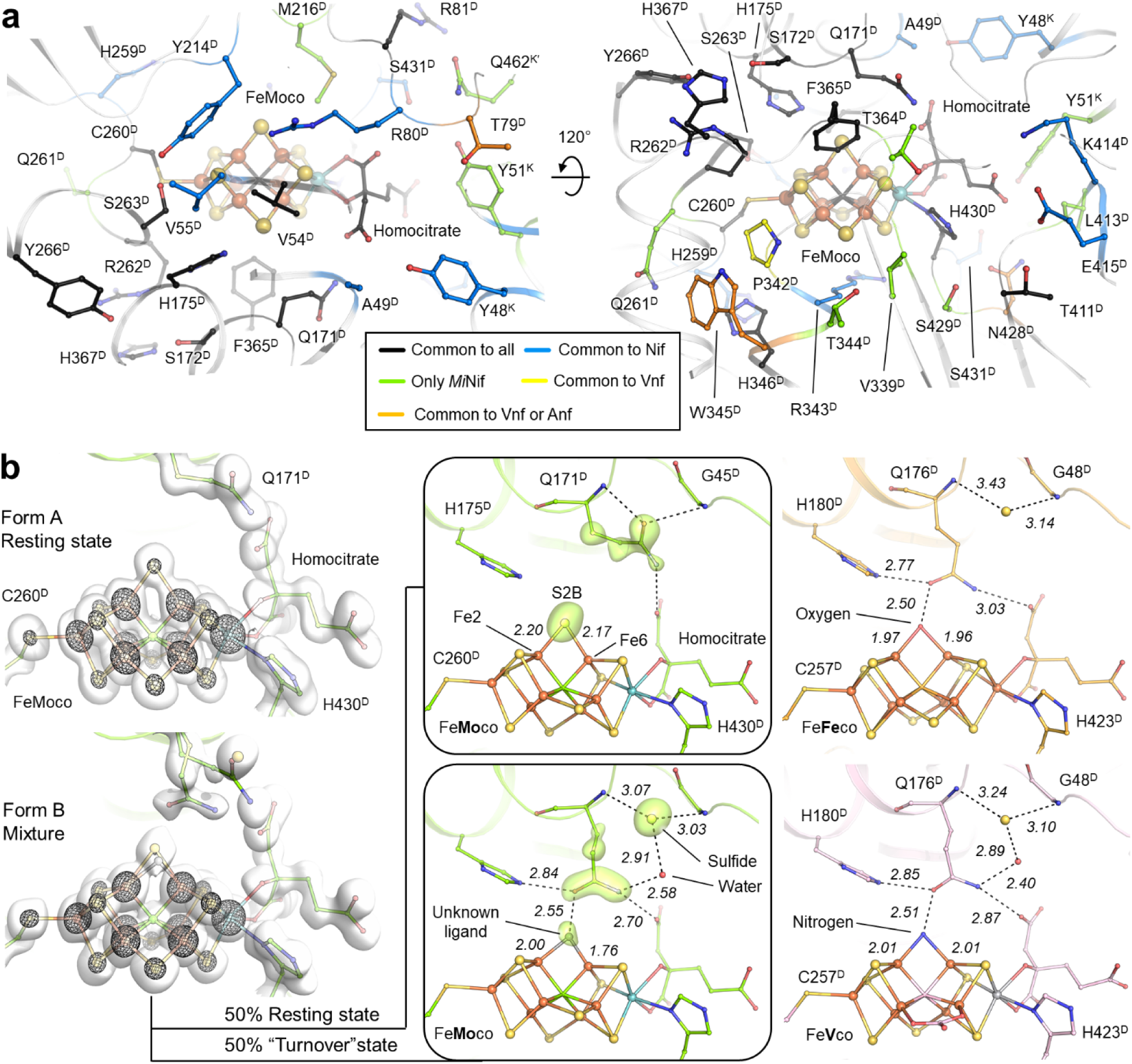
N_2_-reduction site. **a**, Conservation of the FeMoco surroundings in *Mi*NifDK and structural homologues. The protein backbone is shown as cartoons, with residues which are part of the 1^st^ and 2^nd^ spheres displayed as sticks. Metallocofactors are displayed as spheres and sticks. A colour coding of carbon shows the conservation among *Av*NifKD, *Av*VnfDK and *Av*AnfDK. **b**, Close up of the FeMoco in the form A (top) and form B (bottom). Fe and S from the metallocofactor are displayed as spheres, and residues in the close vicinity of the FeMoco are shown as sticks. 2*F*_o_−*F*_c_ maps are overlaid as a transparent surface (1-σ) and black mesh (9-σ). Right panels illustrate the interpretation of the mixture of different states from the form B deconvoluted as resting (top) and turnover (bottom) states. An omit map was generated for Gln171^D^, S2B, and the unknown ligand for the corresponding states, and the resulting difference maps are presented as a green transparent surface contoured to 6-σ. As a comparison, the FeVco/FeFeco site in the turnover states for *Av*VnfDK and *Av*AnfDK is presented on the right. Bond distances are indicated in Å.

The form A harbours the conventional FeMoco in a “resting state” in which the S2B fully occupies its position on the FeMoco sulfur belt and the Gln171^D^ side chain binds the homocitrate and amide backbone Gly45^D^ (Fig. 4). Remarkably, the form B exhibits a mixture modelled with a 50/50 occupancy of the resting state and the so-called “turnover” state that has never been observed in a Mo-dependent nitrogenase (Fig. 4b). The turnover state was discovered in *Av*VnfDK, in which the protein preparation was obtained by diminishing the concentration of reducing agents during the cultivation (47, 60). The same mixture of states was later observed in *Av*AnfDK (42). This state is defined by (*i*) the migration of the S2B from the sulfur belt to the position occupied by the oxo group of the Gln171^D^, (*ii*) a ligand bridging the Fe2 and Fe6, *(iii)* a repositioning of the canonical Gln171^D^ side chain binding the ligand and homocitrate, and *(iv)* a water molecule bridging the Gln171^D^ side chain and the S2B. It is worth noting that the observed movements differ from the CO or selenide-trapped structure, where the conserved glutamine was unable to rotate and sulfide was not present (61, 62). While the readjustments are the same in *Mi*NifDK, the monoatomic ligand is not equidistant between Fe2 and Fe6 as modelled in *Av*VnfDK (PDB 6FEA) and *Av*AnfDK (PDB 8BOQ) and has a shorter distance from the Fe6 (Fig. 4b). The short distance (1.76 Å) is reminiscent of Fe chemical complexes engaging a double bond with a nitrogen while remaining too large for a triple bond (63, 64). Despite the near atomic resolution, the mixture of states prevented us from providing the atom identity of the ligand as we did for the central carbide, and future studies including spectroscopy will have to be conducted to clarify the FeMoco status.

## Discussion

The development of methods for sustainable NH_3_ production to meet the constantly increasing demand through optimisation of N_2_ reduction technologies has become the new frontier of our modern era (7, 28, 65–67). Exploiting the microbial genetic diversity of diazotrophs is a promising strategy for finding robust and more efficient nitrogenases with peculiar properties. For instance, *Mi*NifH and the enzyme involved in the FeMoco biosynthesis, *Mi*NifB, have been in the spotlight for successful transplantation *in planta* (26, 27, 68). The *M. infernus* system is of particular interest for this type of biotechnological application because of its robustness and the minimal *nif* operon, reduced to four structural genes, two regulators, and three possible accessory proteins (30). The native purification performed in this work detected only the presence of the regulator NifI_1,2_ associated with the nitrogenase, suggesting that accessory proteins are not bound to *Mi*NifDK and might have a role in optimising the metallocofactor biosynthesis and enzyme maturation (30). The growth dependency on molybdenum, with a required minimum of 100 nM MoO_4_ (Fig. 1b), alongside a remarkable tolerance to tungstate, showcases an adapted metal trafficking and recognition system that avoids tungsten misincorporation in the nitrogenase cofactor (29). Notably, the work of Payne and colleagues (18) showed that the minimal concentration of molybdenum necessary for diazotrophic growth in *M. infernus* may be drastically lower in its physiological environment, in which molybdenum is supposed to be rare.

Diazotrophic *Methanococcales* living in extreme environments might harbour a relic of the nitrogenase predecessor (20–22). This theory, based on phylogeny and the evolutionary history of methanogens, has been recently reinforced by the discovery that the reactivation system of the methanogenic enzyme (i.e., Methyl-Coenzyme M reductase) contains a metallocluster similar to an intermediate in the FeMoco biosynthesis (69). Altogether, the fact that the biosynthesis of the catalyst for methane generation (i.e., CfbCD) requires a NifDK-like machinery reinforces the scenario that methanogens adapted these enzymes to create the primordial nitrogenase (21, 22). Our structural studies corroborate this hypothesis by highlighting structural simplifications and resemblances between *Mi*NifDK and homologues belonging to NifDK, VnfDK, and AnfDK (Fig. 2 and Supplementary Figs. 3-5), thereby suggesting that the ancestral versions of these homologues were more similar to nitrogenases from *Methanococcales*. Furthermore, the structural insights reveal the perfect conservation of metallocofactor coordination, plausible proton shuttling to the FeMoco centre and the close environment of the S2B, while some parts involved in the second metal coordination sphere, the hydrated cavity surrounding the homocitrate, and the electron transfer path could be more divergent.

*Mi*NifDK natural hyperthermostability and rigidity (Fig. 1e, Supplementary Figs. 5-7), together with the naturally reduced state of the cytoplasm in the methanogen, as well as the rapid purification and crystallisation under strict exclusion of O_2_ and dithionite, could explain why the form B retained the P^1+^ and the partial FeMoco turnover states. Trapping the P^1+^ offers a picture of the transient, electron-depleted physiological state provoked by NifH binding (12, 56). Obtaining the turnover state in a FeMoco-containing nitrogenase confirms the conservation of the described behaviour across all N_2_-fixing enzymes. The turnover state has been postulated to illustrate a FeMoco loaded with nitrogen-species intermediates located between the Fe2 and Fe6 following the displacement of S2B (50), further stabilised by the canonical glutamine sidechain: a snapshot of the nitrogenase in action. However, this reaction mechanism is still challenged (70, 71), and the intrinsic characteristics of *Mi*NifDK described in this work are expected to deepen our understanding of the nitrogenase mechanism in future studies. For instance, its hyperthermostability features that hold high potential for biotechnology could also be employed to isolate new catalytic intermediates through cold trapping and characterisation by atomic-resolution crystallography and spectroscopy, providing invaluable insights for mimicry-based chemistry and translating this knowledge into the bio-inspired design of new alternatives to the current Haber-Bosch catalysts.

## Methods

### Bacterial strains and growth conditions

*Mi*NifDK and *Mi*NifH were purified natively from *Methanocaldococcus infernus* DSM 11812 (Leibniz Institute DSMZ - German Collection of Microorganisms and Cell Cultures, Braunschweig, Germany). The growth medium was prepared according to Maslać et al. 2022 (29) with the following modifications: Substitution of Na_2_SeO_4_ with Na_2_SeO_3_·5H_2_O (2 µM final); the Na_2_WO_4_·H_2_O concentration was increased to 100 µM final, addition of 22 mM Na_2_SO_4_, and pH adjustment to 6.5. Adaptation to diazotrophic conditions, growth, harvesting, and storage of the cells were handled as described in Maslać et al. 2024 (41) with 10 µM of Na_2_MoO_4_.

*M. infernus* was routinely cultivated anaerobically in 30-60 mL medium in 1 L pressure-resistant Duran bottles with 0.25 to 2 mM final concentration of Na_2_S as a reductant and sulfur source, with a preference of 0.5 mM Na_2_S final in molybdate-containing cultures. After vacuuming the headspace of the bottle to −1 bar, it was filled up to 0.5 bar H_2_/CO_2_ (80:20%) and complemented up to 1 bar with 100% N_2_. The inoculum was added in a 1:20 ratio. The cultures were incubated standing without agitation at 75 °C for two days. After roughly 24 h, the gas phase was exchanged by vacuuming (−1 bar) followed by filling up to 0.5 bar H_2_/CO_2_ (80:20%) and completing to 1 bar with 100% N_2_ gas. Cells were harvested at room temperature in an anaerobic chamber containing a gas mixture of N_2_/CO_2_ (90:10) via centrifugation at 6,000 x *g* at 20 °C for 30 min. The wet cell pellet was transferred to a sealed serum flask, 0.8 bar of N_2_ was added, and the flask was stored at −80 °C. All proteins studied in this work come from batch cultures.

*M. infernus* could also be grown in a fermenter, as presented in Figure 1c. Here, biomass production of *M. infernus* was carried out in a 10 L fermenter (BIOSTAT® B plus, Sartorius, Göttingen, Germany) using 7 L of the same culture medium. Anoxic conditions were achieved by sparging the fermenter with N_2_ before inoculation for a few hours. Pre-cultures grown as described above were used as inoculum at a 1:20 ratio. The fermenter was maintained at 75 °C, and 300 μL of Antifoam 204 (Sigma Aldrich, St. Louis, USA) was added immediately before inoculation to prevent foaming. The redox balance and sulfur sources were maintained by supplying a 1.6 M stock of Na_2_S solution continuously, with an initial flow rate of 1.5 mL.min^-1^, which was increased to 3 mL.min^-1^ after 11 h. The culture was continuously sparged with H_2_/CO_2_ (80:20%) at 2 L.min^-1^ and N_2_ at 0.5 L.min^-1^, with stirring at 500 rpm. After 11 h, the gas supply was increased to 5 L.min^-1^ for H_2_/CO_2_ and 1 L.min^-1^ for N_2_, while the stirring speed was raised to 600 rpm. The cultivation continued for 6.5 h and was stopped at an OD_600nm_ of ≈ 0.89. *M. infernus* cells grown in the fermenter were harvested under anaerobic conditions by using an adapted 2 L glassware to collect the cells and transfer them to the anaerobic chamber containing a gas atmosphere of N_2_/CO_2_ (90:10%). The cells were centrifuged at 6,000 x g at 20 °C for 20 min. Pellets were transferred to serum flasks, supplemented with 0.8 bar of H_2_/CO_2_ (80:20%) and stored at −80 °C.

For the physiological experiments testing different temperatures, vanadium addition, molybdate concentrations, and N_2_ pressure, a culture volume of 10 mL was used in 1 L pressure-resistant Duran bottles. Regarding molybdate depletion, several transfers without molybdenum with a limited amount of NH_4_Cl (22 mM final) were made, and the cultures were switched to diazotrophy with insufficient leftover ammonium to promote growth. In this case, molybdate was excluded from the trace elements and added anaerobically directly to the medium. The volume of the inoculum was 1:10 except for the N_2_-dependent growth on Supplementary Fig. 2b, where it was 1:20. The same temperature or medium composition was utilised except for the mentioned varying conditions. Experiments were performed in triplicate with a final sodium sulfide concentration of 0.25-0.5 mM. For the N_2_-dependent growth experiment, different volumes of 100% N_2_ were injected in a 1 L bottle containing 1 bar of H_2_/CO_2_: 0.5 mL (0.02 mmol), 5 mL (0.2 mmol), or 50 mL (2 mmol). The bottle contained 20.2 mmol N_2_ when 0.5 bar of N_2_ was present in the bottle as described above. All values were calculated under standard conditions (i.e., 25 °C). Notably, the moles of N_2_ displayed represents the total quantity in the bottles, not how much N_2_ would be dissolved at 75 °C. The cultures were incubated for 48 hours and were sampled five times.

### Protein purification

The *Mi*NifDK and *Mi*NifH were purified several times with similar protocols. A representative purification protocol is described below. The purification procedure of both proteins was controlled by SDS PAGE, absorbance at 280, 415 and 550 nm, and by visual check of the brown colour of the proteins.

An 11.8 g wet-weight of cells was processed. Frozen cell pellets were transferred to an anaerobic chamber containing a gas atmosphere of N_2_/CO_2_ (90:10%) at room temperature. 60 mL of lysis buffer (50 mM Tris/HCl pH 8.0, 2 mM dithiothreitol (DTT)) was added to lyse the cells through an osmotic shock. The viscous lysate was sonicated using a KE76 probe Bandelin SONOPULS (Sigma, Berlin, Germany) at 75% power for four times, 30 seconds each. The resulting lysate was centrifuged at 45,000 x *g* for 45 minutes at 20 °C. After centrifugation, the supernatant was transferred to a Coy tent containing a gas atmosphere of N_2_/H_2_ (97:3%) under yellow light and filtered through a 0.2 μm filter (Sartorius, Göttingen, Germany). Filtered supernatant was loaded onto a 5 mL HiTrap^TM^ Q Sepharose High-Performance column (GE Healthcare Life Sciences, Munich, Germany), pre-equilibrated in lysis buffer at 3 mL.min^-1^. Proteins were eluted with a 0.2 to 0.5 M NaCl linear gradient in lysis buffer in 45 minutes with a 1 mL.min^-1^ rate (9 column volumes, CV). *Mi*NifDKI_1,2_ eluted mainly between 0.41 to 0.47 M NaCl. Pooled fractions were diluted 1:1 with lysis buffer supplemented with 2 M (NH_4_)_2_SO_4_ and filtered again with a 0.2 μm filter (Sartorius, Göttingen, Germany). The filtrate was loaded onto a Source^TM^ 15PHE 4.6/100 PE column (Cytiva, Freiburg, Germany) pre-equilibrated with lysis buffer supplemented with 2 M (NH_4_)_2_SO_4_ to perform hydrophobic interaction chromatography. After washing the column, proteins were eluted with a 1 M to 0.34 M (NH_4_)_2_SO_4_ linear gradient for 15 minutes at a 1 mL.min^-1^ flow rate (8.8 CV). Fractions containing *Mi*NifDKI_1,2_ eluted between 0.74 to 0.34 M (NH_4_)_2_SO_4_. Pooled fractions were concentrated to 5 mL using a 50-kDa cut-off Vivaspin centrifugal concentrator (Sartorius, Göttingen, Germany). The concentrate was diluted with 25 mL lysis buffer containing freshly prepared 10 mM 2-oxoglutarate, 2 mM MgCl_2_, and 2 mM ATP and incubated overnight inside the Coy tent to separate *Mi*NifDK from *Mi*NifI_1,2_. The sample was filtered through a 0.2 μm filter and loaded onto a 5 mL HiTrap^TM^ Q Sepharose High-Performance column (GE Healthcare Life Sciences, Munich, Germany) pre-equilibrated with the same lysis buffer containing the ligands. After washing, the protein was eluted with a 0.15 to 0.4 M NaCl linear gradient in the same ligand-supplemented buffer in 20 minutes at a 1 mL.min^-1^ flow rate (4 CV). Fractions containing only *Mi*NifDK eluted between 0.34 and 0.39 M NaCl. The sample was then concentrated with a 50-kDa cut-off Vivaspin centrifugal concentrator, and the buffer was exchanged by ultrafiltration for the storage buffer (25 mM Tris/HCl pH 8.0, 10% v/v glycerol, and 2 mM DTT).

*Mi*NifH purification was carried out with the same cell preparation. *Mi*NifH mainly eluted between 0.36 and 0.4 M NaCl during the first anionic exchange chromatography step. This pool was diluted with three volumes of the lysis buffer supplemented with 2 M (NH_4_)_2_SO_4_, filtered through a 0.2 μm filter and loaded on a 5 mL Phenyl Sepharose^TM^ High-Performance column (GE Healthcare Life Sciences, Munich, Germany) pre-equilibrated with the same buffer. After washing, proteins were eluted with a 1.5 to 0.5 M (NH_4_)_2_SO_4_ linear gradient for 75 minutes with a 1 mL.min^-1^ flow rate (15 CV). *Mi*NifH eluted between 1.1 and 0.84 M (NH_4_)_2_SO_4_. Fractions were pooled, and the buffer was exchanged for lysis buffer by ultrafiltration on a 30-kDa cut-off Vivaspin centrifugal concentrator (Sartorius, Göttingen, Germany). The resulting sample was filtered and injected on a 5 mL HiTrap^TM^ Q Sepharose High-Performance column pre-equilibrated with lysis buffer. After washing, the elution was performed by applying a 0 to 75 % linear gradient of 500 mM sodium malate pH 5.4 and 2 mM DTT for 25 minutes with a 1.5 mL.min^-1^ flow rate (7.5 CV). *Mi*NifH eluted between 50 to 62 % of 500 mM sodium malate pH 5.4, and 2 mM DTT. The pooled sample was concentrated with a 30-kDa cut-off Vivaspin centrifugal concentrator (Sartorius, Göttingen, Germany), and the buffer was exchanged for the storage buffer.

### Protein denaturation experiment

To measure the melting point temperature of as-isolated *Mi*NifDK, a denaturation experiment was performed in the exact same conditions as those of the Supplementary Fig. 3 of Maslać et al., 2024, where the assay is downscaled to a 20 µL volume reaction (41). The experiment was performed in duplicate in a Coy anaerobic chamber (Coy Laboratory Products) containing a N_2_/H_2_ gas atmosphere (97:3%). *Mi*NifDK was diluted to a final concentration of 5 mg.mL^-1^ in 50 mM Tris/HCl pH 8.0, 100 mM NaCl, and 2 mM DTT in a volume of 20 μL. Eppendorf tubes containing *Mi*NifDK were incubated for 25 min at different temperatures in a block heater. After incubation, the tubes were immediately centrifuged for 25 min at 6,000 x *g* at 20 °C. 5 μL of each supernatant was diluted 10 times with 50 mM Tris/HCl pH 8.0 and 2 mM DTT before estimation of the protein concentration by the Bradford method, performed in aerobic conditions and using bovine serum albumin (BSA) as standard. The values reported in the graphs of Fig. 1e correspond to the protein quantification ratios of the supernatants before and after heat treatment. The obtained values were compared in Fig. 1e to the values obtained for *Av*NifDK in the work of Maslać et al., 2024 (41).

### UV/Visible spectroscopy

As-isolated *Mi*NifDK was transferred inside an anaerobic chamber filled with a 100% N_2_ atmosphere. Absorbance spectra were recorded at room temperature in a FLUOstar Omega Microplate reader (BMG Labtech, Germany) in 384-well plates. The spectra were measured using 15 µL of 0.3 mg.mL^-1^ *Mi*NifDK prepared in storage buffer, supplemented or not with 5 µL of a 0.5 mM dithionite solution (0.225 mg.mL^-1^ *Mi*NifDK and 0.125 mM dithionite final). Blanks were measured on the storage buffer or the storage buffer supplemented with 0.125 mM dithionite. Absorbance values for both samples were blank-subtracted, and values obtained for *Mi*NifDK with 0.125 mM dithionite were rationalised by multiplying values by a 1.333 factor.

### Activity measurements

Activity assays were carried out as described in Maslać et al. 2024 (41) with a few adjustments in volume, time, and temperature. The final reaction volume was 100 µL, incubated for 30 minutes at 50 °C. Experiments were done in an anaerobic glovebox filled with a gas phase of 100% N_2_. A freshly prepared sodium dithionite solution was used as an electron donor to the system. A *Mi*NifDK:*Mi*NifH molar ratio of 1:16 was used. The reaction mixture contained 100 mM MOPS buffer at pH 7.0, 30 mM phosphocreatine, 5 mM ATP, 0.6 mg.mL^-1^ BSA, 0.2 mg.mL^-1^ creatine phosphokinase (from Rabbit muscle, Merck/Roche, Darmstadt, Germany), 10 mM MgCl_2_, 10 mM dithionite, 0.1 mg.mL^-1^ *Mi*NifDK and 0.5 mg.mL^-1^ *Mi*NifH. All reactants and proteins were prepared in 100 mM MOPS buffer pH 7.0. Traces of (NH_4_)_2_SO_4_ coming from the *Mi*NifH purification generate a background that was subtracted to calculate *Mi*NifDK specific activity (Fig. 1f).

For the *Mi*NifDKI_1,2_ response to 2-oxoglutarate, the pool that eluted from the Source^TM^ 15PHE 4.6/100 PE column was further purified. Two protocols can be used to enrich the protein. In the first one, the pool was diluted with three volumes of lysis buffer, filtered through a 0.2 μm filter and loaded on a 5 mL HiTrap Q Sepharose^TM^ High-Performance column (GE Healthcare Life Sciences, Munich, Germany) pre-equilibrated with the same buffer. After washing, the elution was performed by applying a 0 to 75 % linear gradient of 500 mM sodium malate, pH 5.4 and 2 mM DTT, resulting in the enrichment of *Mi*NifDKI12 in fractions collected between 50 to 62 % of the gradient. In the second protocol, the pool from the Source^TM^ 15PHE 4.6/100 PE column was further purified was supplemented with 1 M (NH_4_)_2_SO_4_ final, filtered through a 0.2 μm filter and loaded on a 5 mL Phenyl Sepharose^TM^ High-Performance column (GE Healthcare Life Sciences, Munich, Germany) pre-equilibrated with the lysis buffer supplemented with 1 M (NH_4_)_2_SO_4_. A gradient from 1 to 0 M (NH_4_)_2_SO_4_ was performed, resulting in the enrichment of *Mi*NifDKI_1,2_ in fractions collected between 0.54 and 0.34 M of the gradient. The pooled sample was then concentrated through a 30-kDa cut-off Vivaspin centrifugal concentrator (Sartorius, Göttingen, Germany) and injected on a Superose 6 increase 10/300 (GE Healthcare Life Sciences, Munich, Germany) after removal of particles through centrifugation or filtering. The elution was performed at a flow rate of 0.4 mL.min^-1^ in the storage buffer supplemented with 150 mM NaCl. The pooled sample was finally concentrated through a 30-kDa cut-off Vivaspin centrifugal concentrator.

The assay was performed with a final amount of 0.008 mg *Mi*NifDK per assay accounting for a molar ratio of 1 mol *Mi*NifDKI_1,2_:0.8 mol of *Mi*NifDK. The activity presented in the Supplementary Fig. 2c is relative to the highest activity measured when the complex recovers full activation at 10 mM 2-oxoglutarate.

Since no reactions were measurable at room temperature in the glovebox (18-20 °C, Fig. 1f), we defined the start of the reaction when the reaction mixtures were placed on a thermal heater. After an incubation at 50 °C for 30 minutes, the reaction was quenched using 30 µL of 400 mM EDTA (adjusted to pH 8.0). NH_3_ measurement was carried out aerobically with the o-phthalaldehyde method (described below, (72–74)), directly or after storage for an overnight at 4 °C. We previously determined that under these conditions, *Mi*NifDKH activity is linear in the first 40 min. Omission of one component/substrate/cofactor was used as a negative control, and three replicates were used for each condition.

NH_3_ measurement was carried out using a solution of 100 mL 0.2 M phosphate buffer pH 7.0 containing 2.7 mg.mL^-1^ o-phthalaldehyde (OPA), 3.5 mM 2-mercaptoethanol and 2% (v/v) ethanol. This reagent mixture was prepared in the dark, where OPA was first dissolved in 5 mL of pure ethanol, and then added to the phosphate buffer solution alongside the 2-mercaptoethanol. 25 µL of standard or sample was added to 1 mL of the OPA reagent mixture. A brief vortexing step was used to homogenise the samples in Eppendorf tubes. 300 µL of the mixture was transferred into a black 96-well plate (FALCON, Germany) in 3 replicates (3 x 300 µL). The plate was covered with aluminium foil and incubated in the dark for 30 minutes. Afterwards, fluorescence was measured using excitation at 410 nm and by measuring emission at 472 nm. Different concentrations of NH_4_Cl solutions were used as standards.

### Crystallisation and structure determination

After purification, samples were centrifuged at 13,000 x *g* for 3 min to remove macro-aggregates and dust, and crystallised inside an anoxic chamber filled with a N_2_/H_2_ (97:3%) atmosphere and maintained at 20 °C. Crystallisation was done by the sitting drop method in 96-well MRC 2-drop polystyrene crystallisation plates (SWISSCI). The crystallisation drop was formed by spotting 0.7 μL of purified protein with 0.7 μL of precipitant. Both crystalline forms were obtained using the same crystallisation condition made of 30 % v/v 2-methyl-2,4-pentanediol, 100 mM Tris pH 8.5, 500 mM Sodium chloride and 8 % w/v Polyethylene glycol 8,000 (crystallisation solution of the JBScreen Wizard form Jena Bioscience, Germany).

*Mi*NifDK crystals form A were obtained with a protein solution at a final concentration of 2.95 mg.mL^-1^. *Mi*NifDK crystals form B were obtained by the same method and crystallisation solution, except that the protein was crystallised at 22.4 mg.mL^-1^.

Crystals of form A took about a month to appear, while the ones from form B grew after a few days and were harvested in a short time following their appearance. Both crystals were directly flash-frozen in liquid nitrogen.

### Data collection and structural analysis

The diffraction experiments used for the deposited model were performed at 100 K on beamlines X06-SA PXI at the Swiss Light Source synchrotron (SLS) for form A and BM07-FIP2 at the European Synchrotron Radiation Facility (ESRF) for form B. The data were processed and scaled with *autoPROC* (75). The data presented a slight anisotropy and hence we used the *STARANISO* correction integrated with the *autoPROC* pipeline (76) (*STARANISO*. Cambridge, United Kingdom: Global Phasing Ltd.). Datasets of form B collected at 19.9 and 20.1 keV were processed without *STARANISO* correction.

The structure of form A was solved by using a computationally generated AlphaFold 2 model as a template of the *Mi*NifDK complex for molecular replacement (77). The structure of form B was solved by using the model of form A as a template for molecular replacement. In both cases, molecular replacement was done with *PHASER* from the *Phenix* package (78). Models were manually built via *Coot* (79) and refined with *phenix.refine* (80). The refinement steps were performed by considering all atoms as anisotropic with hydrogens in the riding position. Models were validated by the MolProbity server (81, 82). Data collection and refinement statistics for the deposited models are listed in Supplementary Table 1. All figures with structures were generated and rendered with PyMOL (Version 2.2.0, Schrödinger, LLC, New York, NY, United States).

A total of four datasets were used to generate the maps in Fig. 2a. The grey-blue density map highlighting the molybdenum atom was created by first generating anomalous maps at 19.9 (Mo K-edge remote) and 20.1 keV (Mo K-edge peak) (Supplementary Table 1). A difference map was generated corresponding to the anomalous map at 20.1 keV, subtracted from the one at 19.9 keV. The signal was finally exemplified by generating a non-crystallographic symmetry (NCS) map from this difference map. The same procedure was applied for the Fe, in which the difference map resulted from subtracting the anomalous map generated at 7.16 keV (Fe K-edge peak) from the one generated at 6.00 keV (Fe K-edge remote). Note that no signals can be visible for the third metal site, in which magnesium was modelled based on its adequate B-factor.

Sequence and structural comparisons to generate graphs of Figures 2a, 2c, and figures of Supplementary Figs. 3, 4, and 5 were done with the following models: *A. vinelandii* NifDK (*Av*Nif) PDB code 3U7Q (54), *A. vinelandii* VnfDK (*Av*Vnf) PDB code 5N6Y (47), *A. vinelandii* AnfDK (*Av*Anf) PDB code 8BOQ (42), *Klebsiella pneumoniae* NifDK (*Kp*Nif) PDB code 1QH8 (53), *Clostridium pasteurianum* NifDK (*Cp*Nif) PDB code 4WES (83), *Gluconacetobacter diazotrophicus* PA1 5 NifDK (*Gd*Nif) PDB code 5KOH (55), and *Rhodobacter capsulatus* AnfDK (*Rc*Anf) PDB code 8OIE (51).

In Fig. 2b, the sequence alignment superposed to the secondary structure of *Mi*NifDK is presented in the Supplementary Figs. 3 and 4 were made with the Espript server (84).

## Data Availability

*Mi*NifDK structures were validated and deposited in the Protein Data Bank (PDB) under the following accession numbers: 9SPZ, Crystal structure of the nitrogenase from *Methanocaldococcus infernus* refined to 1.37 Å resolution, form A. 9SQ3, Crystal structure of the nitrogenase from *Methanocaldococcus infernus* refined to 1.21 Å resolution, form B.

## Author contributions

Organism cultivation, enzyme purification, crystallisation and X-ray data collection were done by all authors. P.B. and N.M established cultivation, diazotrophic adaptation, and performed the different growth experiments. P.B. established and performed the fermenter growth experiment. P.B., N.M., and T.W. established the protein purification and crystallisation. N.M. and M.R.T. established the activity assay. M.R.T. measured enzymatic activities. N.M., P.B., and T.W. performed the data processing, model building, and structure refinement. T.W. validated and analysed the structures and produced the final figures. T.W. wrote the paper with contributions and final approval of all co-authors.

---

**Supplementary Table 1.** Data collection and refinement statistics.

|  | *MiNifDK*<br>form A<br>12.658 keV | *MiNifDK*<br>form A<br>6.000 keV | *MiNifDK*<br>form A<br>7.160 keV | *MiNifDK*<br>form B<br>12.398 keV | *MiNifDK*<br>form B<br>19.900 keV | *MiNifDK*<br>form B<br>20.100 keV |
| --- | --- | --- | --- | --- | --- | --- |
| **Data collection** |  |  |  |  |  |  |
| Wavelength (Å) | 1.00000 | 2.06639 | 1.73160 | 0.97951 | 0.62304 | 0.61684 |
| Space group | *P*2<sub>1</sub> | *P*2<sub>1</sub> | *P*2<sub>1</sub> | *P*1 | *P*1 | *P*1 |
| Resolution (Å) | 78.85 – 1.37<br>(1.52 – 1.37) | 78.99 – 2.24<br>(2.50 – 2.24) | 78.94 – 1.84<br>(2.03 – 1.84) | 101.86 – 1.21<br>(1.35 – 1.21) | 102.00 – 1.81<br>(1.84 – 1.81) | 70.73 – 2.00<br>(2.03 – 2.00) |
| Cell dimensions |  |  |  |  |  |  |
| a, b, c (Å) | 78.43,<br>117.18,<br>106.63 | 78.56,<br>117.72,<br>106.56 | 78.55,<br>117.58,<br>106.56 | 68.50, 80.24,<br>105.74 | 68.60, 80.35,<br>105.89 | 68.54, 80.27,<br>105.80 |
| α, β, γ (°) | 90, 91.48, 90 | 90, 91.58, 90 | 90, 91.60, 90 | 74.49, 82.25,<br>65.02 | 74.49, 82.25,<br>65.02 | 74.50, 82.24,<br>65.0 |
| R<sub>merge</sub> (%)<sup>a</sup> | 9.1 (119.8) | 20.3 (58.0) | 9.8 (53.3) | 7.4 (107.1) | 17.8 (163.3) | 22.2 (167.9) |
| R<sub>pim</sub> (%)<sup>a</sup> | 2.9 (40.4) | 14.5 (47.0) | 6.9 (45.4) | 3.0 (42.5) | 17.8 (163.3) | 22.2 (167.9) |
| CC<sub>1/2</sub><sup>a</sup> | 0.998 (0.658) | 0.984 (0.713) | 0.996 (0.724) | 0.999 (0.669) | 0.980 (0.352) | 0.979 (0.325) |
| I/σ<sub>I</sub><sup>a</sup> | 12.3 (1.9) | 5.6 (1.6) | 8.9 (1.7) | 12.9 (1.7) | 3.5 (0.5) | 2.2 (0.4) |
| Spherical compl. (%)<sup>a</sup> | 68.0 (13.0) | 66.8 (11.9) | 74.0 (14.2) | 62.1 (11.0) | 93.0 (94.1) | 92.5 (92.9) |
| Ellipsoidal compl. (%)<sup>a</sup> | 95.3 (68.2) | 91.8 (61.8) | 93.3 (57.1) | 92.0 (74.5) | / | / |
| Redundancy<sup>a</sup> | 10.4 (9.6) | 5.5 (4.6) | 5.4 (4.3) | 7.1 (7.3) | 3.8 (3.8) | 3.8 (3.8) |
| Nr. unique reflections<sup>a</sup> | 273,586<br>(13,675) | 62,114<br>(3,107) | 124,152<br>(6,209) | 371,908<br>(18,595) | 168,457<br>(8,470) | 122,976<br>(6,176) |
| **Refinement** |  |  |  |  |  |  |
| Resolution (Å) | 21.91 – 1.37 |  |  | 33.62 – 1.21 |  |  |
| Number of reflections | 273,457 |  |  | 371,077 |  |  |
| R<sub>work</sub>/R<sub>free</sub><sup>b</sup> (%) | 12.33/15.56 |  |  | 12.08/14.60 |  |  |
| Number of atoms<sup>c</sup> | 16,851 |  |  | 17,736 |  |  |
| Protein<sup>c</sup> | 15,164 |  |  | 15,422 |  |  |
| Ligands/ions<sup>c</sup> | 233 |  |  | 291 |  |  |
| Solvent<sup>c</sup> | 1,454 |  |  | 2,023 |  |  |
| Mean B-value (Å<sup>2</sup>) | 23.64 |  |  | 17.71 |  |  |
| Molprobity clash<br>score, all atoms | 2.15 |  |  | 2.29 |  |  |
| Ramachandran plot |  |  |  |  |  |  |
| Favoured regions (%) | 96.88 |  |  | 97.15 |  |  |
| Outlier regions (%) | 0 |  |  | 0 |  |  |
| rmsd<sup>d</sup> bond<br>lengths (Å) | 0.011 |  |  | 0.013 |  |  |
| rmsd<sup>d</sup> bond<br>angles (°) | 1.262 |  |  | 1.472 |  |  |
| **PDB ID code** | 9SPZ |  |  | 9SQ3 |  |  |

<sup>a</sup> Values relative to the highest resolution shell are within parentheses. <sup>b</sup> R<sub>free</sub> was calculated as the R<sub>work</sub> for 5 % of the reflections that were not included in the refinement. <sup>c</sup> Hydrogens are excluded from the numbers of atoms. <sup>d</sup> rmsd, root mean square deviation. Compl. Stands for Completeness.

**Supplementary Fig. 1.**
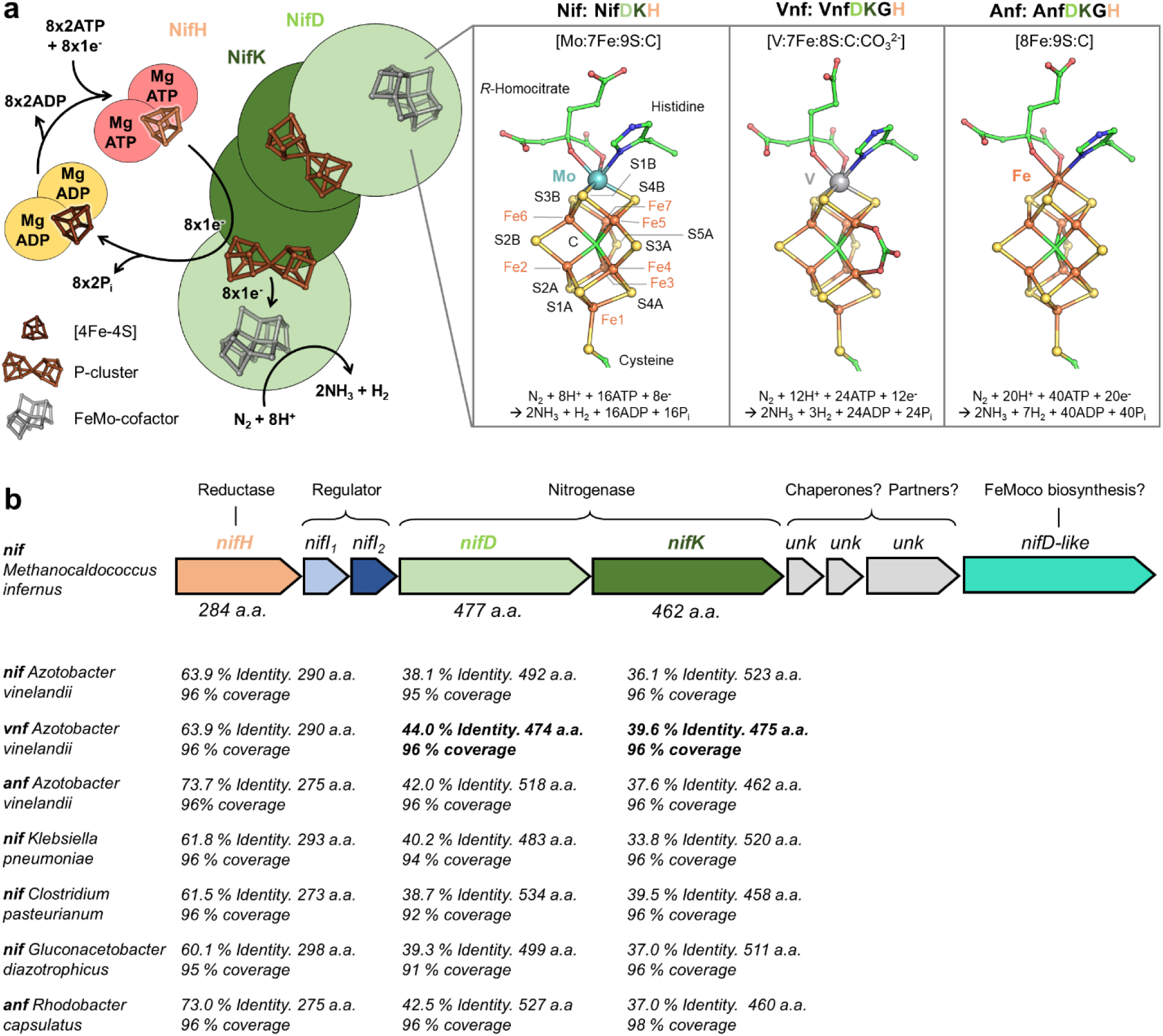
Overview of the nitrogenase cycle and sequence comparison between structural homologues and *M. infernus nif* system. **a,** Presentation of the nitrogenase catalytic cycle in which reduced NifH loaded with MgATP provokes the displacement of an electron from the P-cluster to the FeMoco following the electron-deficient mechanism previously proposed (13). The P-cluster in its P^1+^ state switches back to its fully reduced P^N^ state by uptaking the electron from NifH upon ATP hydrolysis. Phosphate release (P_i_) has been shown to be the rate-limiting step of the cycle (85). Different types of catalysts exist exhibiting different chemical properties (86)(right panel): the FeMoco, FeVco, and FeFeco carried by NifDK, VnfDK, and AnfDK, respectively. The overall reaction model comes from (87) **b**, *nif* operon in *M. infernus* containing three putative genes of unknown function (*unk*, (30)). Amino acid lengths (a.a.) are indicated for the protein encoded by *nifDKH*, and their protein sequence identity/coverage is compared to homologues. In the archaeal kingdom, only methanogens and anaerobic methanothrophs are diazotrophs (88). Therefore, they are not only key actors in the carbon cycle, generating or degrading approximately 1 Gt of methane annually (89), but also contribute to the nitrogen cycle in specific niches (23). *Methanococcales* depend exclusively on H_2_ and CO_2_ (or formate) to generate methane, a catabolism that has been described as at the thermodynamic limits of Life (90). Despite these low energy yields, they can sustain the energetic toll of at least 16 ATP per fixed N_2_ thanks to the unique post-translational regulation system (28) and reallocation of cellular resources (29), which are not shared with the studied diazotrophic bacteria. Note that *nifB*, critical for the FeMoco synthesis, is located at a different position in the genome.

**Supplementary Fig. 2.**
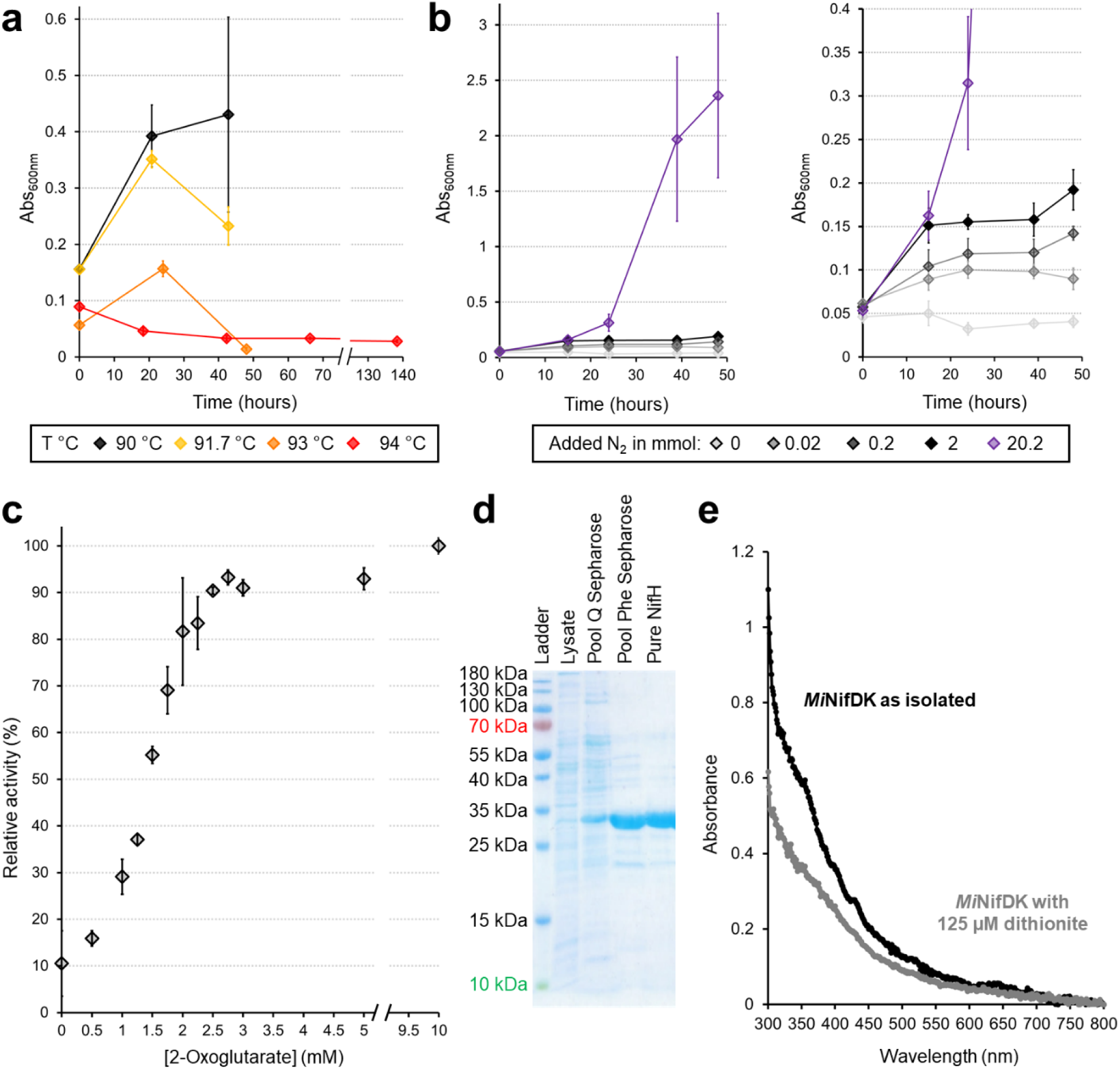
Physiological capability of diazotrophic *M. infernus*, isolation of *Mi*NifH and partially oxidised *Mi*NifDK. **a,** Growth curves of *M. infernus* grown diazotrophically at different temperatures. The slight increase in Abs_600nm_ at 93 °C was not considered as growth. Diazotrophic precultures for the experiment at 93 °C and 94 °C were first adapted to 90 °C before inoculation. **b,** Growth curves of *M. infernus* with different N_2_ amounts. The right panel is a close-up of the low N_2_ addition. **c,** 2-oxoglutarate-dependent inhibition of *Mi*NifI_1,2_ relative to the maximum activity recorded at 10 mM 2-oxoglutarate. The NH_3_-production activity assay was performed by combining *Mi*NifDKI_1,2_ with *Mi*NifH at 50 °C under 100% N_2_ at pH 7.0 for 30 min. A K_0.5_ of 1.425 mM was estimated by testing a 0 to 10 mM 2-oxoglutarate range. A control without ATP and containing 10 mM 2-oxoglutarate did not provide a signal for NH_3_ production (relative activity of 0.1 ± 0.6 compared with added ATP) **d,** Native purification of *Mi*NifH used for the activity assay. In this case, the last chromatography step contains (NH_4_)_2_SO_4_, and despite intense washing through ultrafiltration, the purified NifH fraction was still contaminated with NH_4_^+^, explaining the background observed in Fig. 1f. **e,** UV/visible spectrophotometry profile of as-isolated *Mi*NifDK before and after addition of 125 µM dithionite final.

**Supplementary Fig. 3.**
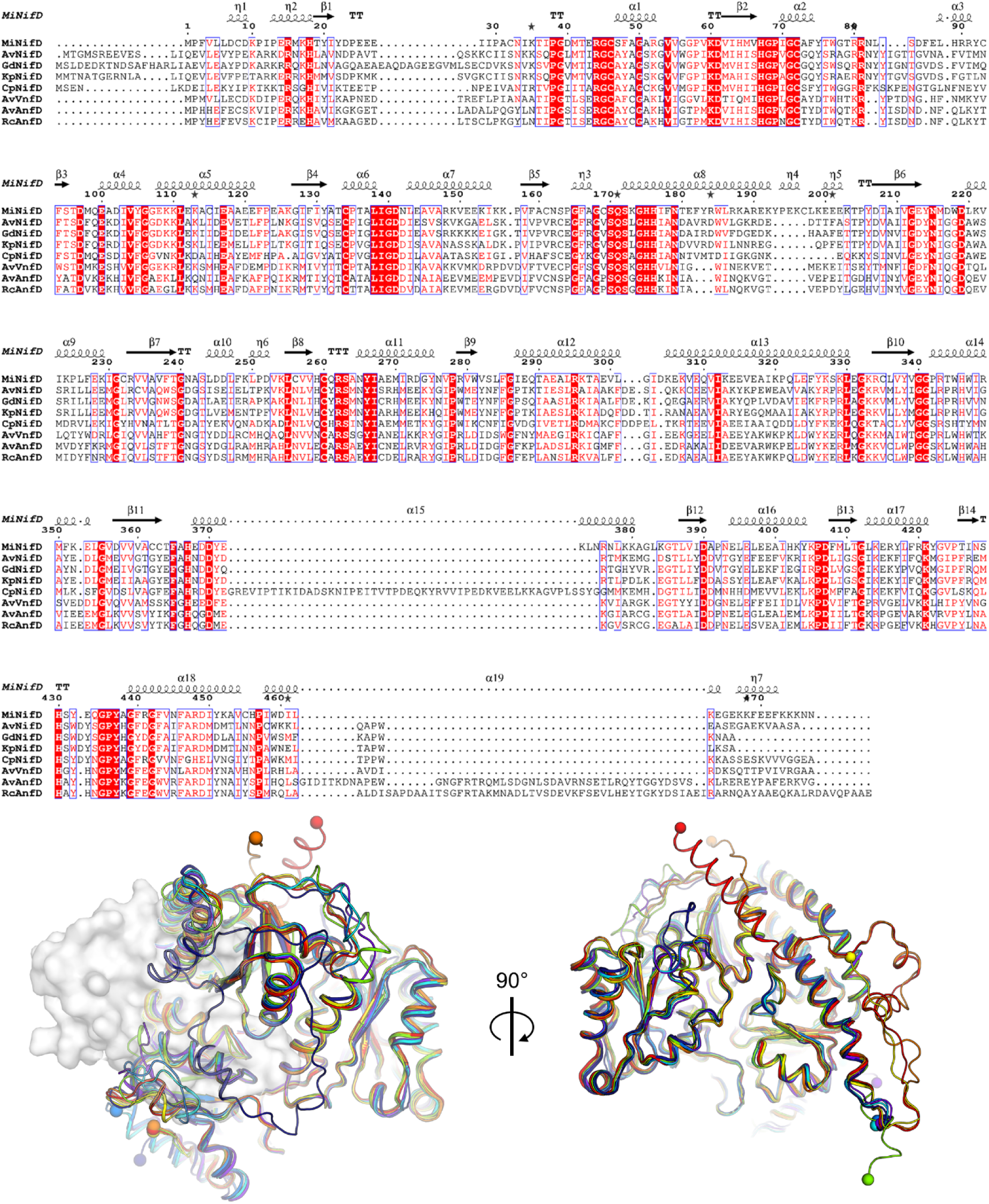
Sequence and structural alignment of *Mi*NifD with homologues. The secondary structure derived from *Mi*NifD with the sequence alignment has been performed with the Espript server (84). The bottom panel represents two views of NifD, with the protein backbone shown as cartoons. The white surface corresponds to *Av*VnfG. Structures are colored as follow: *Mi*NifD form B (green), *Av*NifD (cyan), *Gd*NifD (purple), *Kp*NifD (blue), *Cp*NifD (dark blue), *Av*VnfD (yellow), *Av*AnfD (orange), *Rc*AnfD (red).

**Supplementary Fig. 4.**
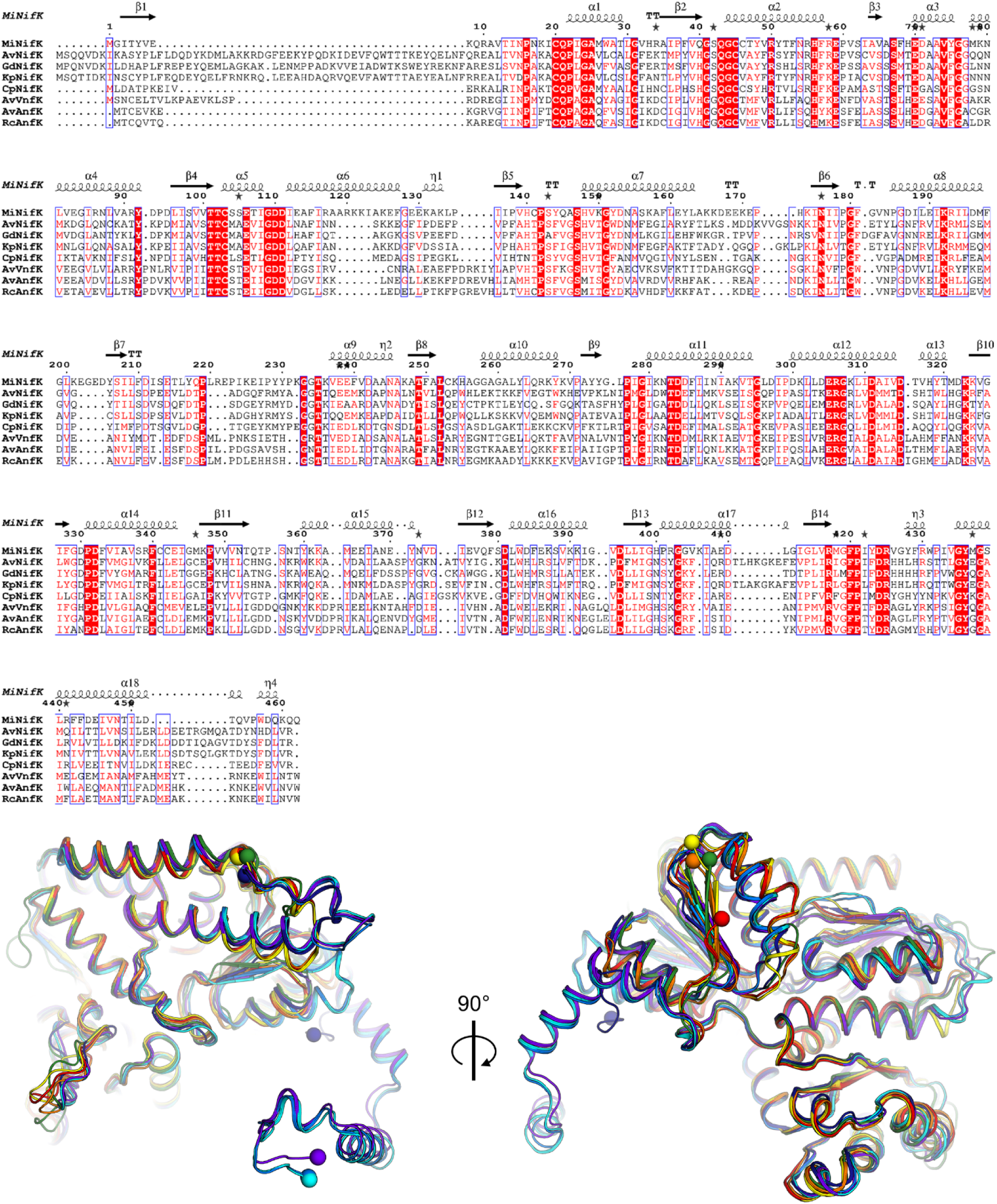
Sequence and structural alignment of *Mi*NifK with homologues. The secondary structure derived from *Mi*NifK with the sequence alignment has been performed with the Espript server (84). The bottom panel represents two views of NifK with the protein backbone shown as cartoons. *Mi*NifK form B (dark green), *Av*NifK (cyan), *Gd*NifK (purple), *Kp*NifK (blue), *Cp*NifK (dark blue), *Av*VnfK (yellow), *Av*AnfK (orange), *Rc*AnfK (red).

**Supplementary Fig. 5.**
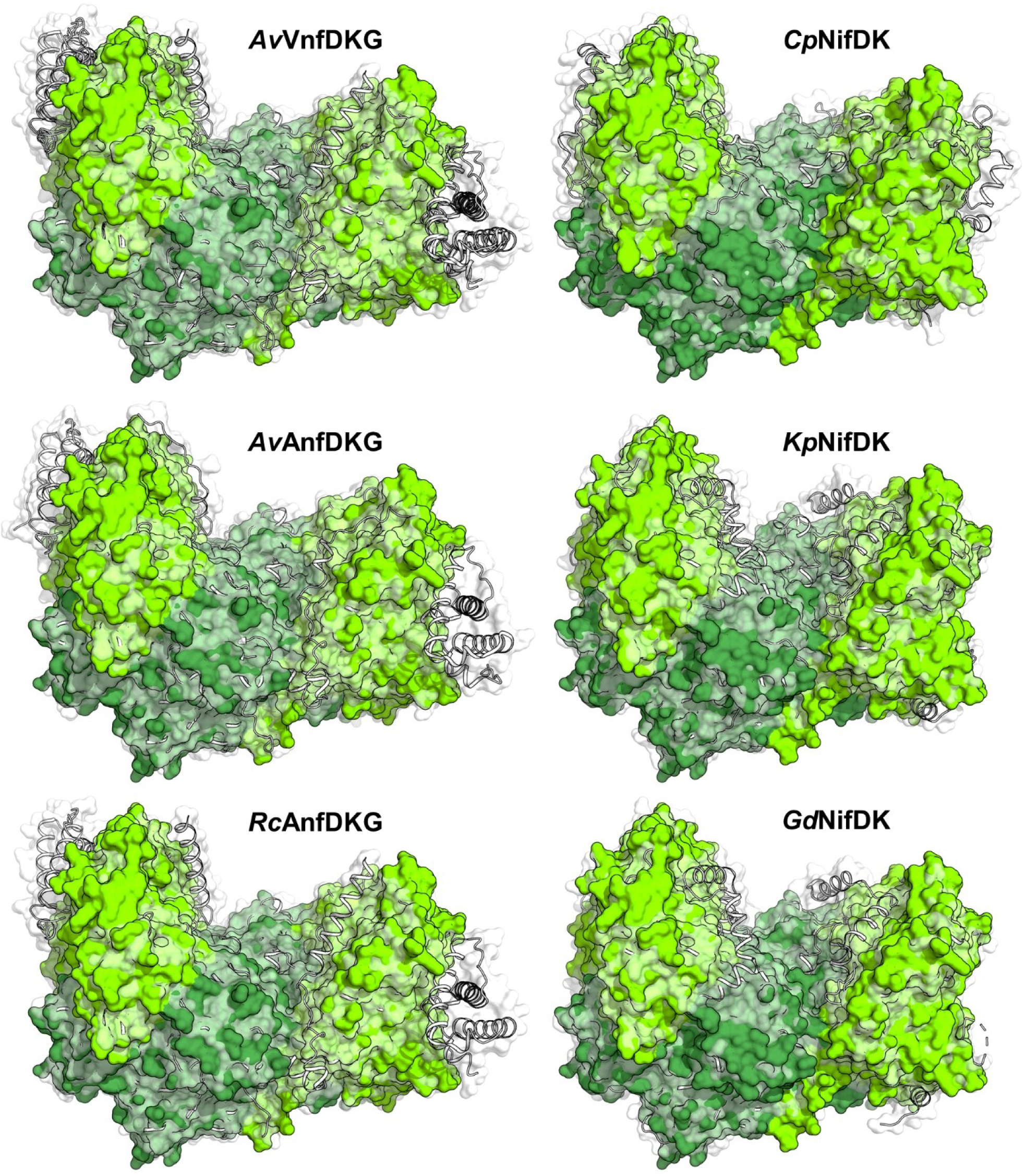
Overall reduction of *Mi*NifDK compared to structural homologues. *Mi*NifD (light green) and *Mi*NifK (dark green) are shown as plain surfaces with superposed structural homologues displayed as a white transparent surface and white cartoons.

**Supplementary Fig. 6.**
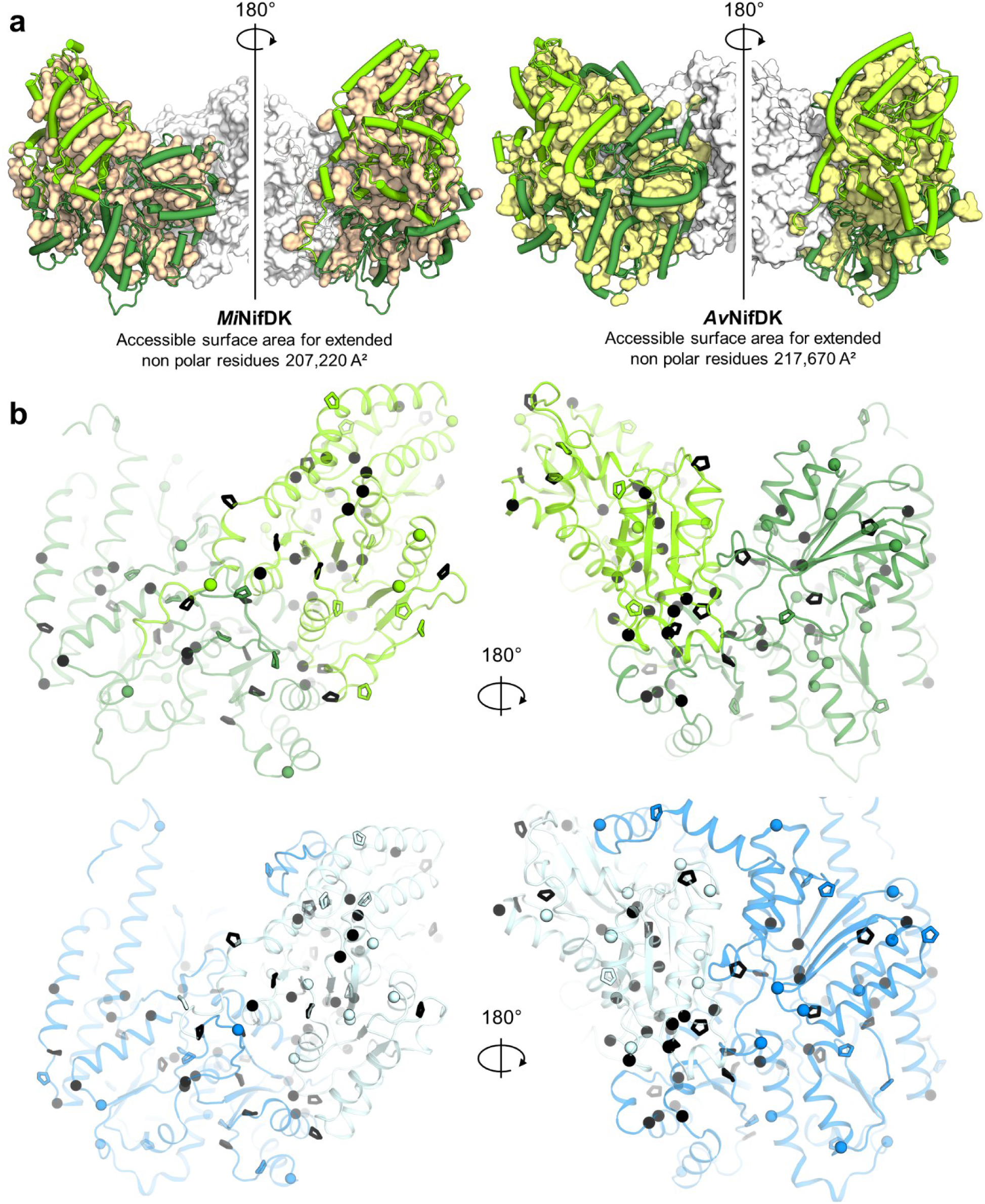
*Mi*NifDK composition in terms of hydrophobic clusters, glycine and proline and comparison to *Av*NifDK. **b,** Hydrophobic clusters (side chains of Val, Leu, Met, and Phe) in *Mi*NifDK (left) and *Av*NifDK (right). One NifDK is shown as a cartoon, and the second one as a white surface. The hydrophobic clusters are shown as a wheat/yellow surface. **c,** *Mi*NifD (light green), *Mi*NifK (dark green), *Av*NifD (cyan) and *Av*NifK (blue) are shown as cartoons with glycine Cα displayed as spheres and proline as sticks. Glycine and proline residues shared between *Mi*NifDK and *Av*NifDK are colored in black.

**Supplementary Fig. 7.**
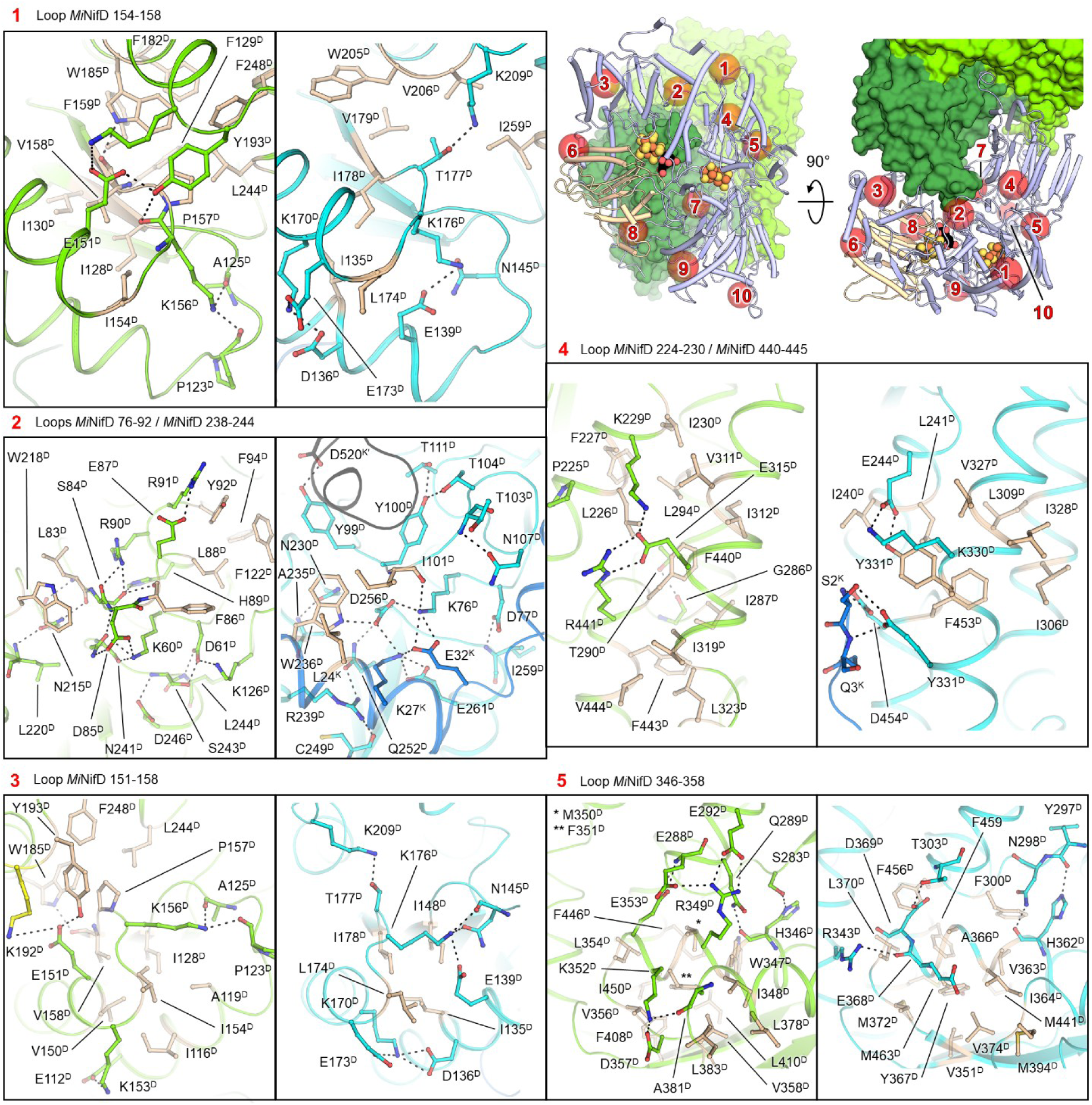
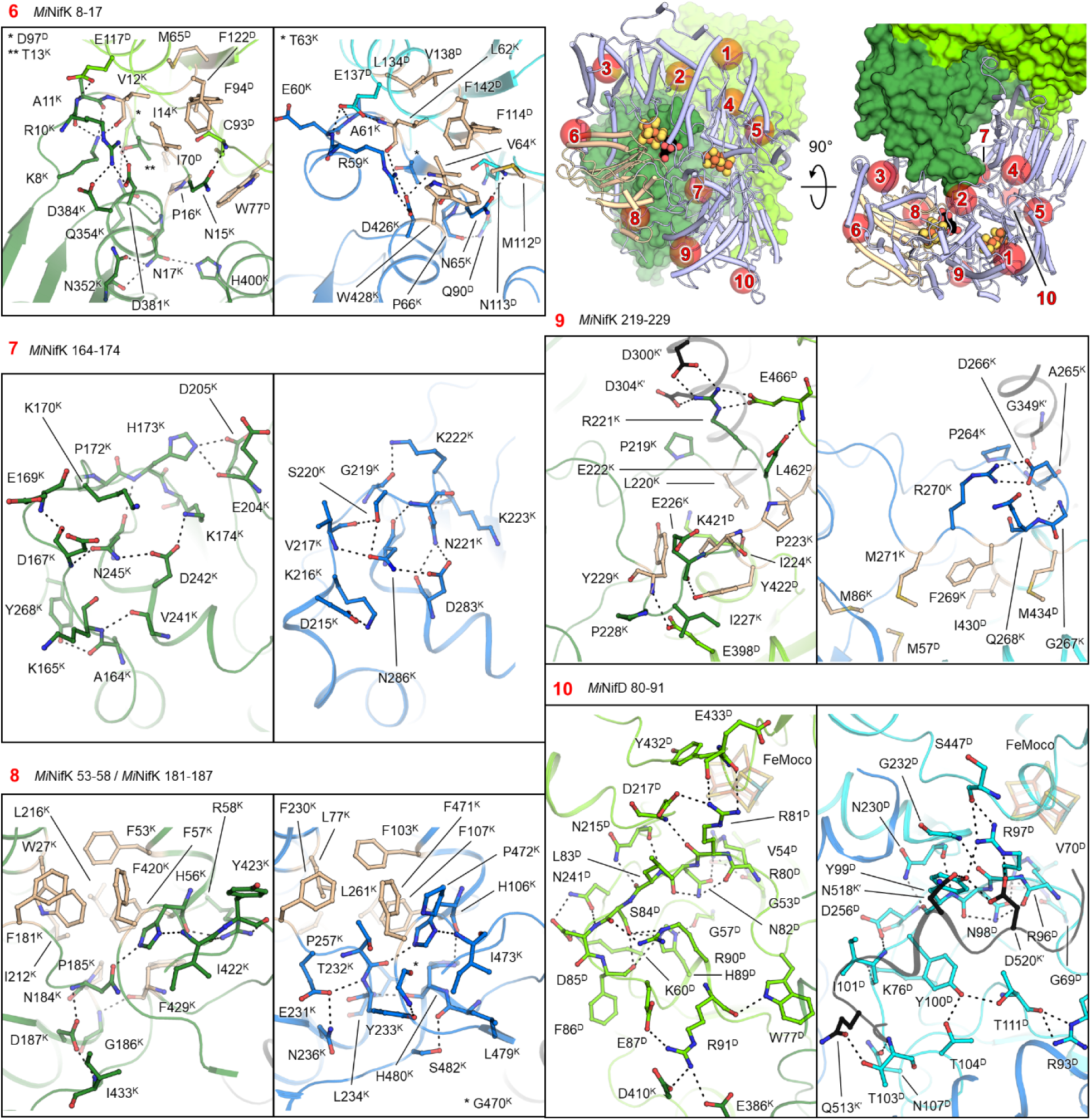
Hot spots in *Mi*NifDK. Top right panel presents the overview of *Mi*NifDK displayed in the same fashion as in Fig. 2c, without the superposition with *Av*NifDK. The cartoon coloured in wheat indicates the αIII domain known for its intrinsic dynamics (50, 51). Local hot spots correspond to a structural reinforcement between *Mi*NifDK and *Av*NifDK through the apparition of salt bridges/hydrogen bonds (highlighted by dashes) or an increase of Van der Waals contacts through hydrophobic side chains (highlighted by wheat sticks). Inserts represent close-ups of the numbered area with the protein backbone shown in cartoon and the residue of interest in sticks or spheres. *Mi*NifD, *Mi*NifK, *Av*NifD, and *Av*NifK are colored green, dark green, cyan, and marine, respectively, in the close-ups. The opposite NifK dimer is colored in black. Top right panel presents the overview of *Mi*NifDK displayed in the same fashion as in Fig. 2c, without the superposition with *Av*NifDK. The cartoon coloured in wheat indicates the αIII domain known for its intrinsic dynamics (50, 51). Local hot spots correspond to a structural reinforcement between *Mi*NifDK and *Av*NifDK through the apparition of salt bridges/hydrogen bonds (highlighted by dashes) or an increase of Van der Waals contacts through hydrophobic side chains (highlighted by wheat sticks). Inserts represent close-ups of the numbered area with the protein backbone shown in cartoon and the residue of interest in sticks or spheres. *Mi*NifD, *Mi*NifK, *Av*NifD, and *Av*NifK are colored green, dark green, cyan, and marine, respectively, in the close-ups. The opposite NifK dimer is colored in black.

**Supplementary Fig. 8.**
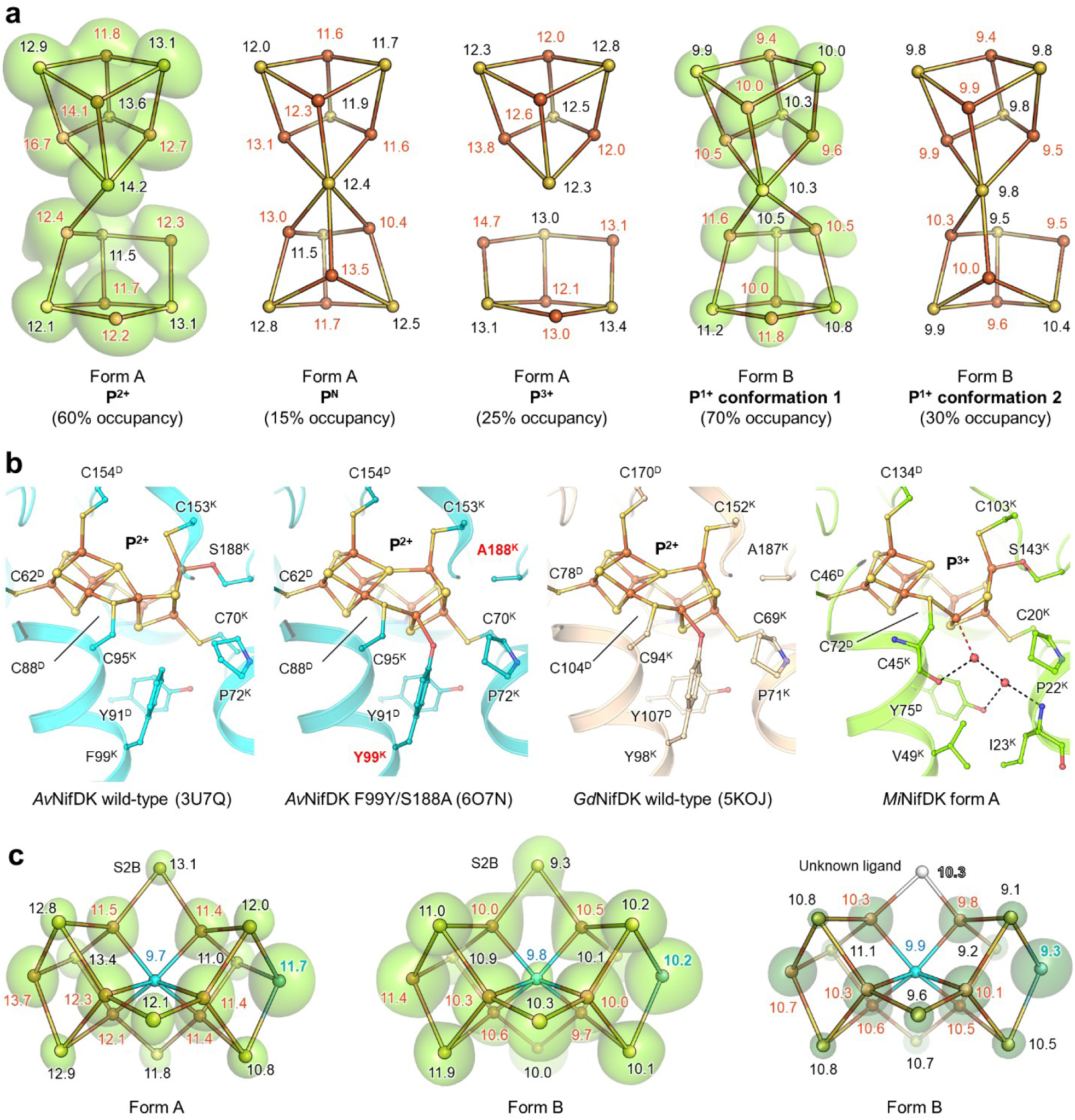
Omit maps for the metallocofactor and comparison of the P-cluster binding in different models. **a**, Omit maps and b-factors of the P-cluster in the two forms. *F*_o_−*F*_c_ maps are presented as a green transparent surface contoured to 5-σ and 15-σ for form A and form B, respectively. The different modelled states are displayed in balls and sticks, and individual atomic b-factors are labelled with S and Fe in black and red, respectively. **b**, Differences in the oxidised P-cluster coordination between AvNifDK wild-type/variant, NifDK from *Gluconacetobacter diazotrophicus* (*Gd*NifDK) and *Mi*NifDK. Residues in the close vicinity of the P-cluster are displayed as balls and sticks, with waters in red spheres. Dashes show wateŕs interactions with the surrounding. The water-Fe8 distance shown in red is 2.7 Å. **c**, Omit maps and b-factors of the FeMoco in the two forms. *F*_o_−*F*_c_ maps are presented as a green transparent surface contoured to 15-σ, 6-σ, and 23-σ for form A, form B with modelled S2B, and form B with modelled ligand, respectively. The different modelled states are displayed in balls and sticks, and individual atomic b-factors are labelled with S and Fe in black and red, respectively. The unknown ligand was refined as a nitrogen.

**Supplementary Fig. 9.**
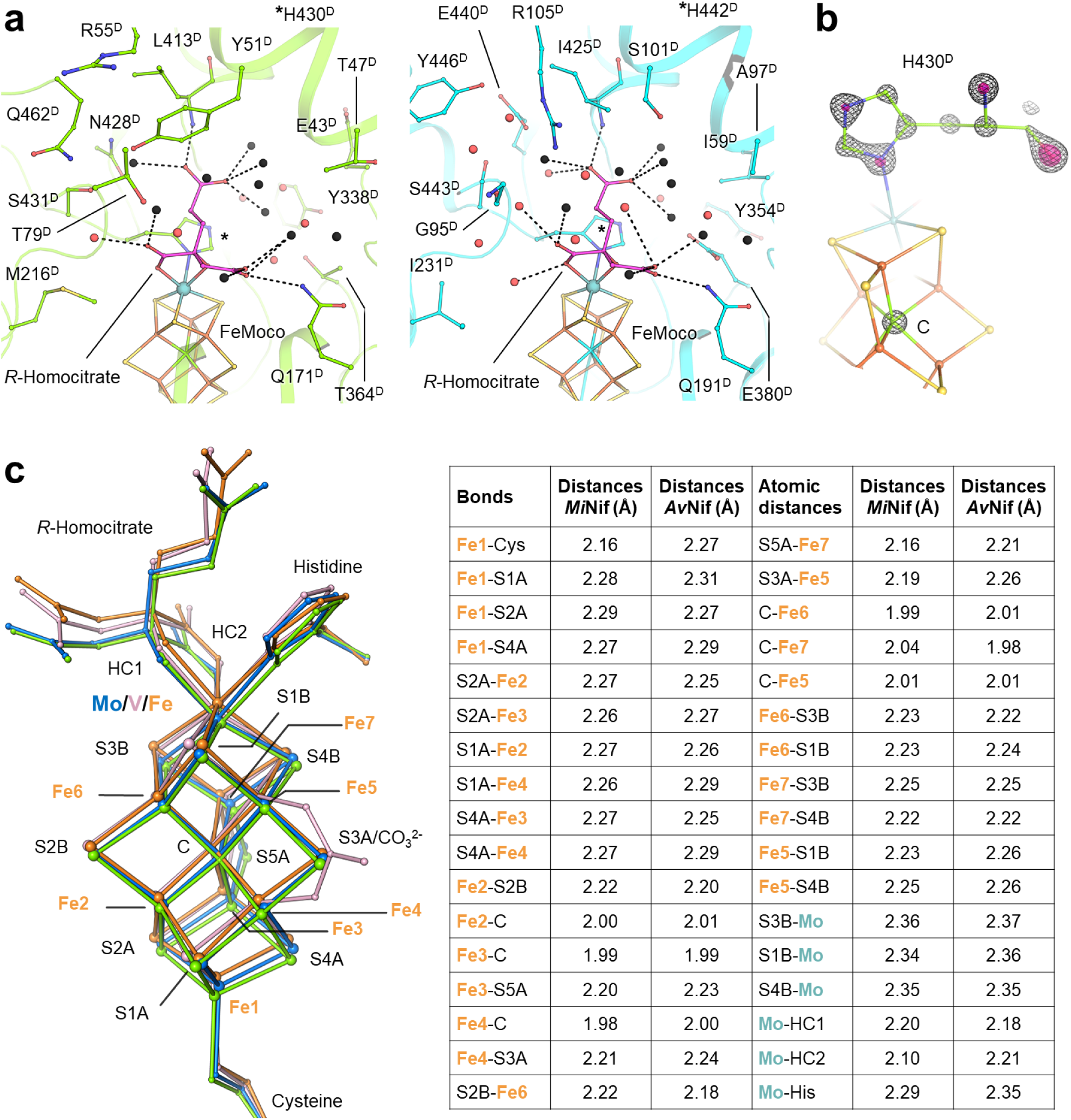
Comparison of the water network surrounding the homocitrate and comparison in the carbide-containing FeMoco. **a.** Water network in *Mi*NifD (green, left) and *Av*NifD (cyan, right) around the *R*-homocitrate (pink). Waters are shown as spheres and coloured black or red if they collocate (with a distance below 0.7 Å) or not between *Mi*NifDK or *Av*NifDK. Residues in the close vicinity and ligands are depicted as balls and sticks. Dashes indicate direct contact between the homocitrate and the water molecules. **b**. Close up of the FeMoco presenting an electron density for the interstitial carbon similar to a carbon rather than a nitrogen or oxygen. The 2*F*_o_−*F*_c_ map is contoured at 4.5 σ (black mesh) and 6.5 σ (transparent pink surface). **c**. Superposition of the FeMoco, VFeco, and FeFeco, from *Mi*NifDK (green), *Av*NifDK (cyan), *Av*VnfDK (pink), and *Av*AnfDK (orange). Superposition and distances were done on chain C.

## Acknowledgements

We thank the Max Planck Institute for Marine Microbiology and the Max Planck Society for their continuous support. We also thank Christina Probian and Ramona Appel for their continuous technical support in the Microbial Metabolism laboratory. We acknowledge the French Biology/Health Panel Review Committee for the provision of synchrotron radiation beamtime at the ESRF (Grenoble, France) on beamline BM07-FIP2. BM07-FIP2 is supported by the French ANR PIA3(France 2030) EquipEx+ project MAGNIFIX under grant agreement ANR-21-ESRE-0011. We thank the SLS for the beamtime allocation and the staff of beamline PXI at the SLS and BM07-FIP2 at the ESRF. We would also like to thank Dr Ross D. Milton, Dr Olivier N. Lemaire, Dr André G. Gouveia, Mélissa Belhamri, and Sylvain Engilberge for their comments and suggestions to improve the manuscript.

